# In vivo lineage tracing across human tissues using methylation barcodes in the protocadherin gene cluster

**DOI:** 10.64898/2026.02.23.707349

**Authors:** Samuel F. Hackett, Christopher T. Boniface, Adriana V.A. Fonseca, Akemi D. Ramos-Yamasaki, Caroline Watson, Hannah M. L. Bazin, Amanda Tan, Henson Lee Yu, Lars L. P. Hanssen, Harveer Dev, Sophia Apostolidou, Aleksandra Gentry-Maharaj, Sadik Esener, Usha Menon, Jamie Blundell

## Abstract

Resolving the lineage history of human cells is fundamental to understanding ageing and cancer but remains hampered by a lack of native, high-resolution markers. Here, we identify the protocadherin (PCDH) gene cluster as a naturally occurring, highly diverse methylation barcode. While PCDH methylation creates neuronal diversity in the brain, we show that stochastic methylation patterns in this region are maintained as heritable, evolvable lineage markers across multiple non-neuronal tissues, including blood, kidney, prostate, and bladder. By tracking these barcodes in serial samples over a decade, we reveal clonal dynamics with high fidelity, quantitatively recapitulating genetic clone sizes. Crucially, PCDH barcodes identify “cryptic” clonal expansions invisible to standard driver-mutation sequencing and resolve subclonal architectures via continuous epimutation. This native barcoding system provides a scalable, driver-agnostic framework for reconstructing somatic evolution in humans.

## Introduction

Lineage tracing—the prospective tracking of cells and their descendants—is a foundational tool in developmental biology ^1^, stem cell research ^2–4^ and cancer ^5,6^. In the past few decades a suite of powerful technologies has been developed for in-vivo lineage tracing, including inducible fluorescent reporters ^7–9^, DNA barcodes ^10–15^, expressible barcodes ^16–18^, transposon-based labelling ^19–21^, and self-editing CRISPR-based systems ^22–24^. While transformative, these engineered approaches have limitations: they are generally not applicable to human tissues, can perturb the system under study, and are often limited in the number of lineages that can be independently labelled. An ideal lineage tracing tool would instead be native to the organism, highly scalable, and possess a vast and evolvable diversity of labels to resolve complex lineage histories in human tissues over time.

Somatic mutations are one such native system. Acquired at a rate of approximately 10^*−*8^ per base pair per cell per year ^25,26^, they create a unique genetic fingerprint for nearly every cell, enabling high-resolution reconstruction of developmental and somatic evolutionary histories. This apsproach has been powerfully applied to study human embryogenesis ^27,28^, childhood cancers ^29^, and the dynamics of both healthy ^30–32^ and malignant tissues ^33^. However, their utility is constrained by practical challenges. The inefficiency of discovering mutations scattered across the genome necessitates costly whole-genome sequencing, which can typically only be applied to single-cell-derived colonies ^30,32,33^, small micro-dissected cell populations, or costly single-cell approaches, all of which have limited ability to scale.

An alternative approach is to use DNA methylation ^34–36^. Epigenetic marks undergo somatic epimutations at rates several orders of magnitude higher than genetic mutations ^35,36^, offering the potential for more targeted and efficient lineage tracing. Recent work has shown that somatic epimutations of specific “fluctuating” CpGs (fCpGs) can be used to detect clonal expansions and track the evolutionary dynamics of cancers ^34–36^. Somatic epimutations have been used for lineage tracing in haematopoiesis but so far these approaches have required single-cell data ^37^. Building on these findings, we hypothesised that certain genomic regions might contain arrays of CpGs that act as natural “methylation barcodes”. In such a system, the pattern of methylation along a single DNA molecule would compress genome-scale lineage information into a compact locus, making it amenable to readout from bulk sequencing in a cost-effective manner.

To test this hypothesis, we performed deep enzymatic methylation sequencing on a cohort of serial pre-leukaemic blood samples for which ground-truth clonal dynamics had previously been established using deep genetic sequencing ^36,38^. Using this data we provide evidence that CpGs in the protocadherin (PCDH) gene cluster function as such a barcode. The PCDH locus encodes a large family of cell-surface recognition molecules which facilitate neuronal self-avoidance by generating combinatorial cell-surface diversity ^39–43^. This combinatorial diversity is generated not by genetic rearrangement, but by the stochastic choice of gene promoters, a process mediated by random DNA methylation ^42,44^. While the PCDH genes are not widely expressed outside the nervous system, we found that the complex methylation patterns established at this locus are sufficiently diverse and heritable to serve as lineage markers across multiple non-neuronal tissues.

## Results

### The PCDH gene cluster is a methylation barcode

To discover genomic loci whose methylation states act as natural lineage markers we analysed a unique collection of serial blood samples collected annually for up to 10 years from 50 women who went on to develop AML and 50 age-matched cancer-free controls ^36,38^ (Fig. 1a). We extracted methylation “barcodes” (binary patterns of methylation at 10 adjacent CpGs along paired reads) from deep targeted enzymatic methylation sequencing (EM-seq) (Supplementary Note 1). The choice of 10 CpGs was a pragmatic trade-off between barcode complexity (favouring longer molecules) and coverage (favouring shorter molecules). Reasoning that in vivo barcodes marking lineages should be diverse in polyclonal samples, we sought genomic loci with highly diverse 10-mer barcode usage across molecules quantified using entropy. Barcodes from the PCDH gene cluster were consistently ranked among the most diverse across our panel (Fig. 1b). The CpGs directly upstream of exons in the PCDH gene cluster are known to undergo stochastic (de)methylation to generate cell-specific *pcdh* isoforms that facilitate self-avoidance in neuronal cells ^39–42^ (Fig. 1c). Our data demonstrates that CpGs across this region undergo extensive stochastic diversification that we reasoned may act as a natural methylation barcode.

**Fig. 1.**
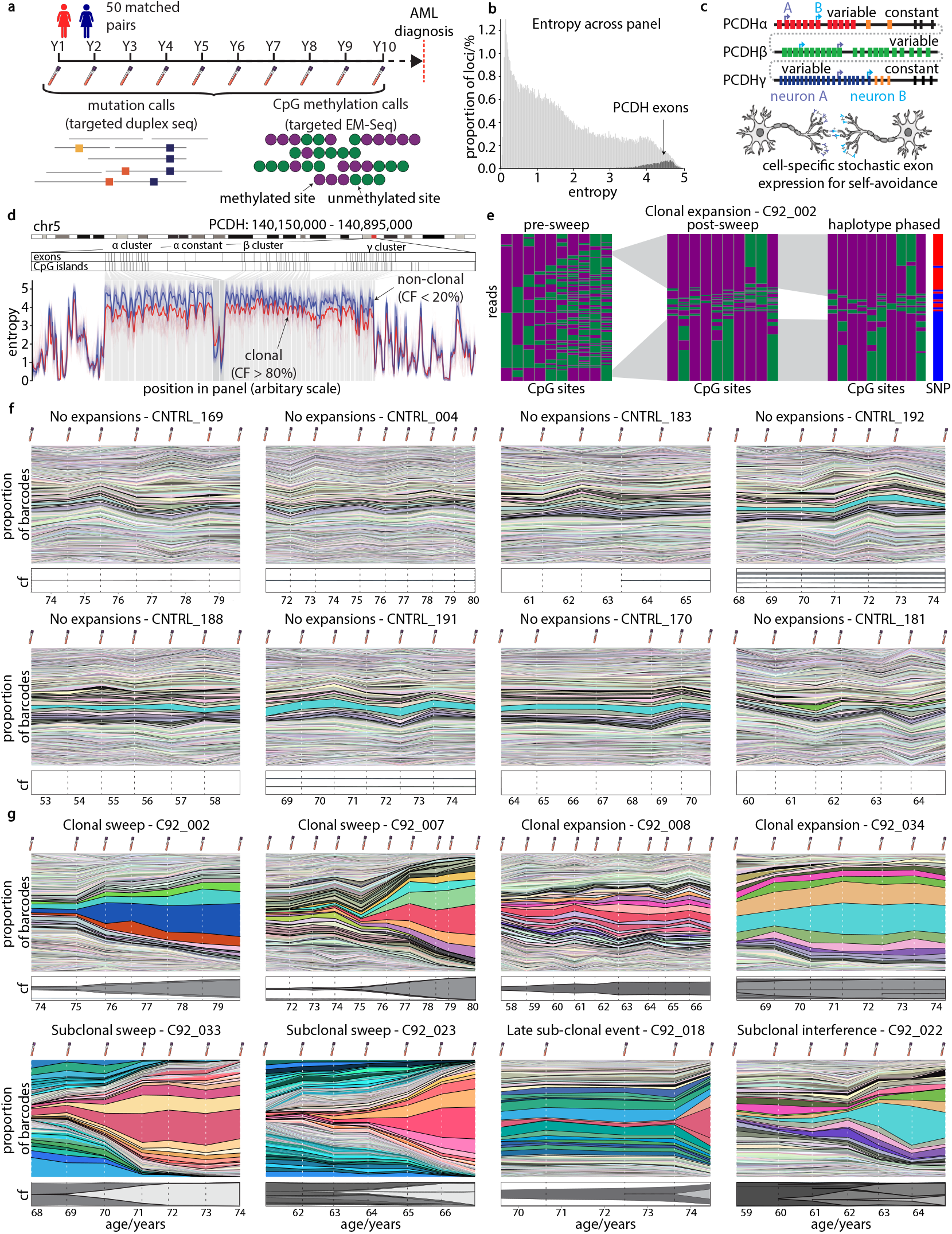
The PCDH cluster serves as a high-entropy native barcode. **a**. Study design: Longitudinal sampling and methylation sequencing of 50 pre-AML and 50 control individuals. **b**. Methylation entropy across the panel, identifying PCDH (arrow) as a locus of extreme diversity. **c**. In neurons, PCDH diversity drives self-avoidance; this study investigates this diversity across other human tissues. **d**. Entropy collapse in clonal samples (individual samples thin red lines, average thick red line) vs. diverse polyclonal controls (individual samples thin blue lines, average thick blue line). **e**. Visualization of barcode ‘sweep’ at 10-mer starting at hg19 chr5:140167837. A diverse methylation landscape (left) is replaced by two dominant haplotypes (right) during clonal expansion. **f**. The barcode-fractions (coloured regions, top panel) measured at the ten-mer with CpG sites in the range hg19 chr5:140256590-140256669 at each time-point for 8 healthy controls. Each coloured region corresponds to a unique epiallele pattern, and the height of these regions corresponds to the read fraction of this pattern at the given time point. The bottom panel shows the somatic cell fraction (cf). **g**. The same for 8 pre-AML donors with large clonal expansions.

To assess the potential of PCDH cluster methylation for clonal lineage tracing, we considered how barcode entropy differed between individuals with known clonal expansions and those without detectable expansions (Fig. 1d). In samples with large clonal expansions (cell fraction > 80%), we observed a significant reduction in the diversity of methylation barcodes across all exons in PCDH excluding the constant domains (Mann-Whitney U test, *p<* 10^*−*2^). This reduction in entropy was driven by the expansion of two sets of closely related barcodes which separate when phased to germline SNPs (Fig. 1e, methods, Supplementary Fig 1). This observation is consistent with a single cell ancestor of the clonal expansion where the two dominant patterns (barcode sets) correspond to the maternal and paternal haplotypes. The patterns which expand at a given locus are different across individuals (Supplementary Note 14), suggesting they do not reflect AML specific reprogramming. The residual diversity within each cluster suggests a process of ongoing epimutation, which we consider in detail later.

Next, we took advantage of our time series data to assess how well PCDH methylation barcodes could be used as clonal lineage markers. We tracked the abundance of each of the 2^10^ = 1024 possible patterns for a 10-mer CpG barcode at a single PCDH locus through time. This revealed a striking difference between pre-AML cases and controls. Polyclonal samples from healthy individuals displayed a highly diverse and stable repertoire of barcodes over many years (Fig. 1f). In contrast, in individuals undergoing a clonal expansion a few barcodes increase in frequency simultaneously with the growth of the somatic clone (Fig. 1g, Supplementary Note 13). These observations show that methylation barcodes at a single locus can be used to efficiently track lineages during somatic evolution.

### Methylation barcodes quantitatively track clone size

We then investigated whether PCDH methylation barcodes could provide quantitative estimates of clone sizes. To test this, we determined the size of the largest somatic variant (SNV or mCA) detected in our targeted panel across serial samples (*n* = 430) for each donor as a ground truth for clone size. We then defined a clonality metric (barcode fraction, BF) as the fraction of the two most common methylation barcodes at a given locus. This metric aims to track clone sizes using the read fractions of the two founding barcodes associated with the maternal and paternal haploid genomes of the single cell ancestor in the expansion. The average barcode fraction across the entire PCDH region closely tracks groundtruth clone size for the majority of samples (Fig. 2a, black points). The observed distribution fits with simulations of a process in which clonal expansion drives the founding barcodes to high frequencies whilst also diversifying via epimutation (Fig. 2a, grey points, methods).

**Fig. 2.**
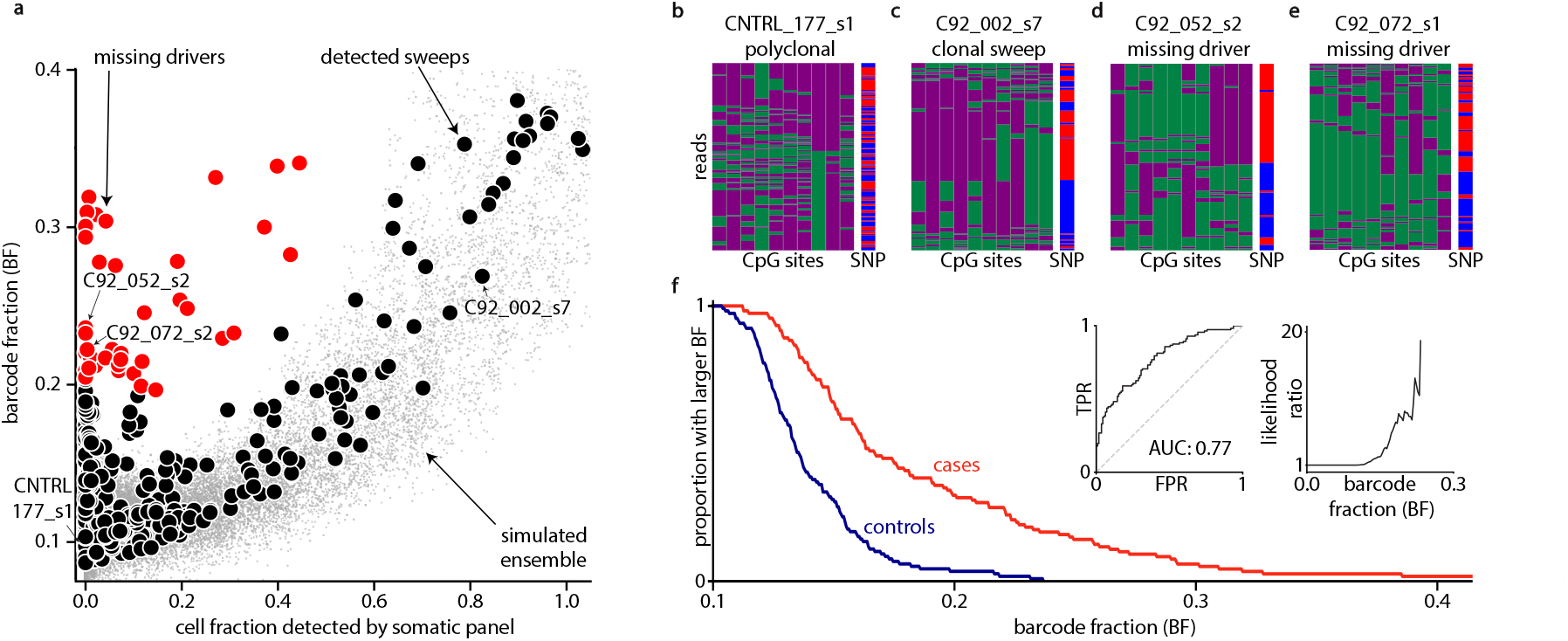
Methylation barcodes allow for driver-agnostic detection of clonal expansions. **a**. Concordance between genetic clone size (x-axis) and methylation barcode fraction (y-axis). Red points indicate samples with high barcode clonality but no detectable genetic drivers, revealing cryptic expansions. **b–e**. Example sets of methylation patterns across samples with no expansion, a detected expansion and two cryptic expansions. **f**. Reverse cumulative curves demonstrating that PCDH barcode fraction is substantially elevated in pre-AML cases relative to controls. Insets - ROC and likelihood ratio at different barcode fractions, showing significant enrichment for pre-AML status.

We reasoned that the quantitative relationship between the barcode fractions and the size of the largest clone would enable us to uncover cryptic clonal expansions; those whose somatic mutations were not detected by the original targeted sequencing panel. In 42 samples (13 pre-AML cases and 3 controls) with no detectable somatic mutations (Fig. 2a red points, methods), methylation patterns bore a close resemblance to those formed in samples with clonal sweeps (Figs. 2b–e), suggesting the presence of a large clonal expansion which was not detected in the mutation panel.

As the presence of expanded clones (high VAF clonal haematopoiesis) is a known risk factor for AML development, we hypothesised that PCDH barcode fraction alone would enable us to identify individuals at higher risk of progression to AML without the need for broader genomic sequencing. To test this we performed deep EM-seq on a further *n* = 112 pre-AML cases and *n* = 111 age-matched controls from the single-time point arm of the original UKC-TOCS study ^45^. The pre-AML cases were sampled at a median of 8 years before AML diagnosis. PCDH barcode fraction is significantly higher in pre-AML cases than in controls (Fig. 2f, Mann-Whitney U test, p *<* 10^*−*6^). Large PCDH barcode fractions strongly enrich for pre-AML cases (insets). Together, these results demonstrate the PCDH methylation barcodes can be used to detect clonal haematopoiesis in a driver-agnostic manner.

### PCDH barcodes diversify and detect subclonal events

The two sets of barcodes that expand with somatic clones during a sweep generally consist of one large barcode and a set of closely related barcodes with lower abundance (Fig. 3a). These observations are consistent with heritable barcodes tracking a clonal expansion whilst undergoing ongoing diversification via de novo epimutations (Supplementary Note 4c). To quantify the epimutation rates, we compared the post-sweep set of barcodes present in the blood samples to a large set of sweeps simulated with a given methylation and demethylation rate (methods). We found that the distribution of barcodes generated during sweeps is consistent with methylation and demethylation rates of ∼10^−2^/year and ∼10^−3^/year, respectively (Fig. 3b). These rates are consistent with values inferred from independent inference frameworks (methods).

**Fig. 3.**
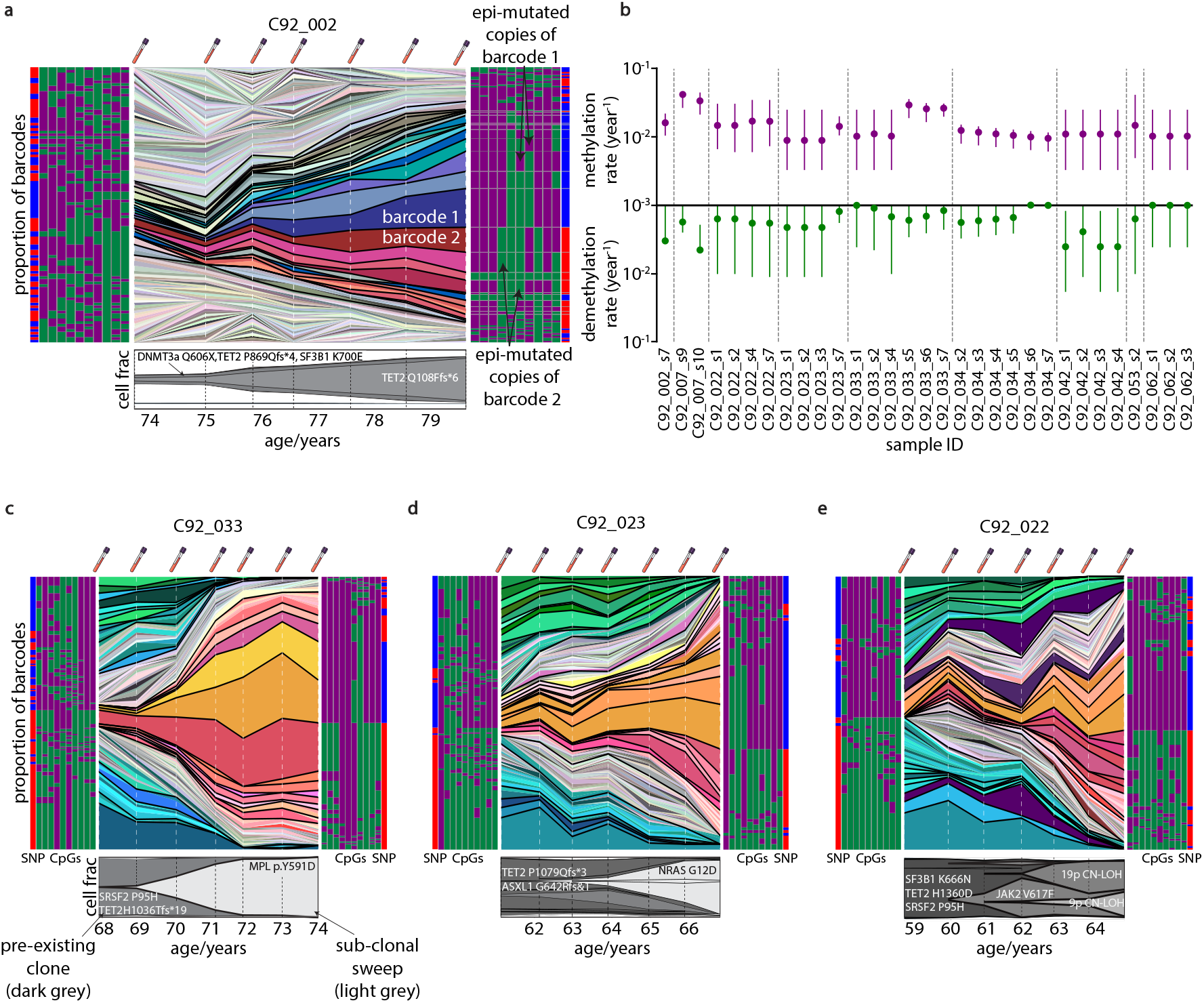
Ongoing epimutation enables the resolution of subclonal dynamics. **a**. Longitudinal dynamics (Muller plot) of PCDH barcode fractions during a clonal sweep. The two founding haplotypes (dark red, dark blue) dominate, while derived epialleles (lighter shades) continuously emerge via de novo epimutation. The pileups (left, right) shows the distribution of these evolved barcodes at the first and last time point. **b**. The estimated methylation (purple, top) and demethylation (green, bottom) rates over samples with large sweeps. Error bars indicate uncertainty due to incomplete sweeps and unknown timing of clonal sweeps. **c**,**d**. Detection of a subclonal sweeps in patient C92_033 and C92_023. The expanding subclone carries a barcode signature derived directly from the ancestral clone, differing by one or two methylation changes. **e**. Clonal interference in patient C92_022, where PCDH barcodes capture the expansion, contraction, and re-expansion of competing subclones (associated with JAK2 and 19p CN-LOH drivers).

The ongoing diversification of PCDH barcodes encodes information about somatic evolution even after a clonal sweep. We previously identified two pre-AML cases with complete sub-sweeps (C92_033 and C92_023). In these donors, somatic mutations show that the first sweep was present prior to initial sample collection, while the second sweep occurred during the sampling period (Fig. 3c,d, bottom).The low diversity of methylation barcodes at early time points reflects the presence of a pre-existing expansion. We then observed the emergence of a sub-sweep reflected in the increased abundance of a subset of barcodes (Fig. 3c,d, Extended Data S17). Importantly, the expanding barcodes associated with the sub-sweep are closely related to the preexisting expanded barcodes. In both donors, the distribution of barcode abundance before and after the sub-sweep suggest that the unobserved first sweeps took substantially longer than the observed second sweeps (Supplementary Note 6), consistent with the later sweeps being significantly more fit as has been previously reported ^38^.

We observed evidence of more complex subclonal competition between JAK2 and 19p CN-LOH subclones in donor C92_022 (Fig. 3e, bottom). Methylation barcodes reflect the subclonal dynamics previously characterized by somatic mutations in which a JAK2-bearing subclone (purple shading) expands between the ages of 60 and 62 years and then is out-competed by the 19p CN-LOH subclone prior to diagnosis.

In principle these complex dynamics are encoded in the relationships between methylation barcodes even from single timepoints. However, due to the low complexity of 10-mer barcodes these more complex subclonal dynamics can only be resolved with time series data. We reasoned, however, that longer barcodes composed of many dozens of informative CpGs might contain sufficient complexity to resolve sub-clonal dynamics from a single time point.

### Long-read PCDH barcodes reflect clonal architecture

To test whether longer methylation barcodes provide sufficient information to infer phylogenetic relationships from bulk sequencing data, we built agent-based simulations to model clonal expansions alongside the dynamics of longer methylation barcodes of various lengths (50-200 CpGs) using realistic epimutation rates. We found that barcodes consisting of *>* 20 CpGs provided sufficient information for lineage reconstruction using traditional phylogenetic inference methods (Supplementary Fig. S7, methods).

To test whether this approach could be used for phylogenetic reconstruction in vivo, we performed long-read whole genome methylation sequencing on peripheral blood DNA from a patient with myeloproliferative neoplasm (MPN, donor “PD6646”) where a high resolution phylogeny was previously built from WGS of single-cell derived colonies ^33^ (simplified in Fig. 4a). We obtained a 280X mean depth whole genome with 10.3kb mean read length and haplotype-phased the aligned reads using germline SNPs (methods). Due to random CpG dropout in long-reads (∼10%-20% per read), we constructed haplotype-specific barcode matrices from sliding windows across the PCDH region that attempted to maximize the number of barcodes (reads) while maintaining mutual coverage of at least 25 CpGs (Fig. 4b and Supplementary Fig. S9, methods, Supplementary Table S1). Hierarchical clustering of each barcode matrix (based on Hamming distance) yielded dendrogram topologies that closely resembled the ground truth phylogenetic tree (Fig. 4c) and dendrograms built from paternal vs maternal haplotypes were highly concordant with each other (Fig. 4c, haploid A, left, and, haploid B, right). We then quantified the fractions of reads assigned to clusters as a proxy for clade size. Remarkably, despite being built from epimutations, barcode matrices recapitulated the known clonal fractions in the ground-truth tree previously built from somatic mutations (Fig. 4d). To check that barcode matrices were specifically reflecting clade sizes we generated long read data from a highly-polyclonal sample as a negative control (donor “KX004”, previous reported by Mitchell *et al*. (2022) ^32^). As expected these data produced clade size inferences that were not consistent with the MPN patient indicating that the barcode inferences in PD6646 were reflecting clade size specific to that patient.

**Fig. 4.**
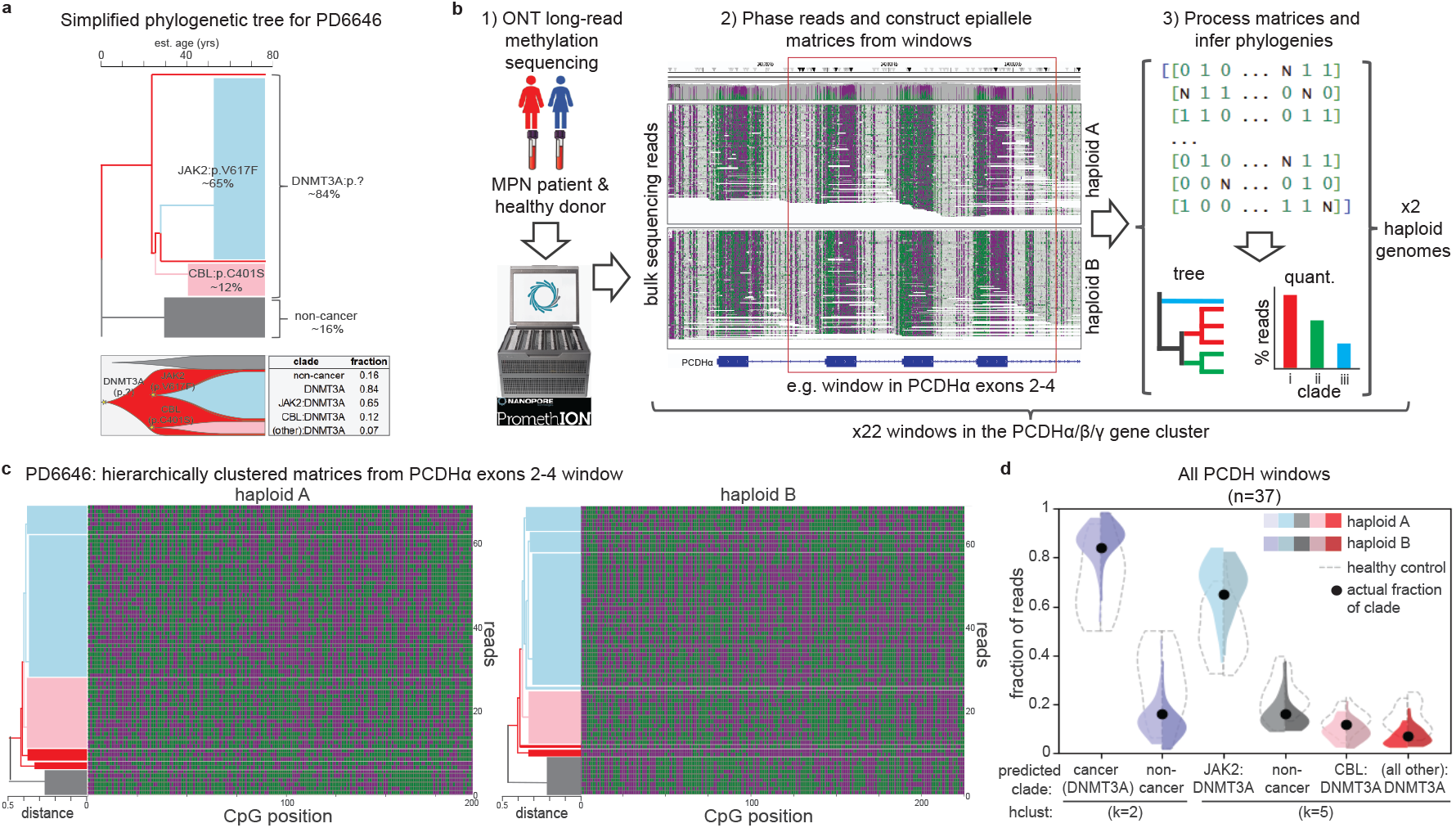
Phylogenetic reconstruction of an MPN patient with long-read epialleles. **a**. Ground-truth phylogeny of an MPN patient (PD6646) derived from single-cell WGS. The clonal architecture features a dominant DNMT3A-mutant clone (∼84%) containing subclonal JAK2, CBL, and other drivers. **b**. Experimental workflow: bulk long-read sequencing (ONT) of peripheral blood is haplotype-phased to generate binary epiallele matrices across the PCDH region which are then clustered by similarity. **c**. Hierarchical clustering of PCDH barcodes from a single window (PCDHa exons 2-4) reconstructs the phylogenetic structure. Coloured bars (left) indicate agreement with the ground-truth clades from (a). **d**. Quantitative accuracy of lineage inference. The distribution of clonal fractions inferred from PCDH barcodes (hierarchical clustering) across all windows (*n* = 37) recapitulates the ground-truth genetic clone sizes (black dots), distinguishing the MPN patient from polyclonal controls (dashed lines).

Methylation barcodes were unable to consistently recapitulate the clade branch lengths found in the ground-truth tree, likely due to the high level of CpG dropout and calling error in the long-read data. Nevertheless, these observations demonstrate that long-read methylation barcodes from the PCDH region can semi-quantitatively resolve the clonal architecture of a sample from a single time point.

### PCDH is an in vivo evolvable barcode across tissues

To investigate whether methylation patterns in the protocad-herin region act as natural barcodes in other human tissues we considered methylation sequencing datasets across a further three tissues: kidney, prostate, and bladder and sought to investigate if the PCDH region alone could provide an assessment of clonality that distinguished between tumours and adjacent normals.

We first investigated a dataset of clear cell renal cell carcinoma (ccRCC) tumour samples (*n* = 122) and adjacent normals (*n* = 63) taken from 68 individuals ^46^. We calculated the average methylation entropy across PCDH in tumours and adjacent normals (Fig. 5a). The PCDH gene cluster displayed a remarkably similar entropy profile to that observed in blood, and showed a highly significant reduction in diversity between tumour (red) and adjacent normal samples (blue) (Mann-Whitney U test, *p<* 10^−6^). To confirm that PCDH methylation barcodes acted as markers of clonality rather than malignancy, we considered a dataset of benign oncocytomas (*n* = 18). These displayed a similar reduction in diversity (orange) as observed in ccRCC samples and bloods with large expanded clones, suggesting barcode diversity collapses due to clonal expansion of founding barcodes (Fig. 5b). Despite the more limited sequencing depth (median 29X) and sample purity (median 55%) methylation barcode fraction across the PCDH region can be used to distinguish tumours from normal tissue with high accuracy (Figure 5c). Barcode fraction also provided a lower bound on tumour purity (Supplementary figure 15b).

**Fig. 5.**
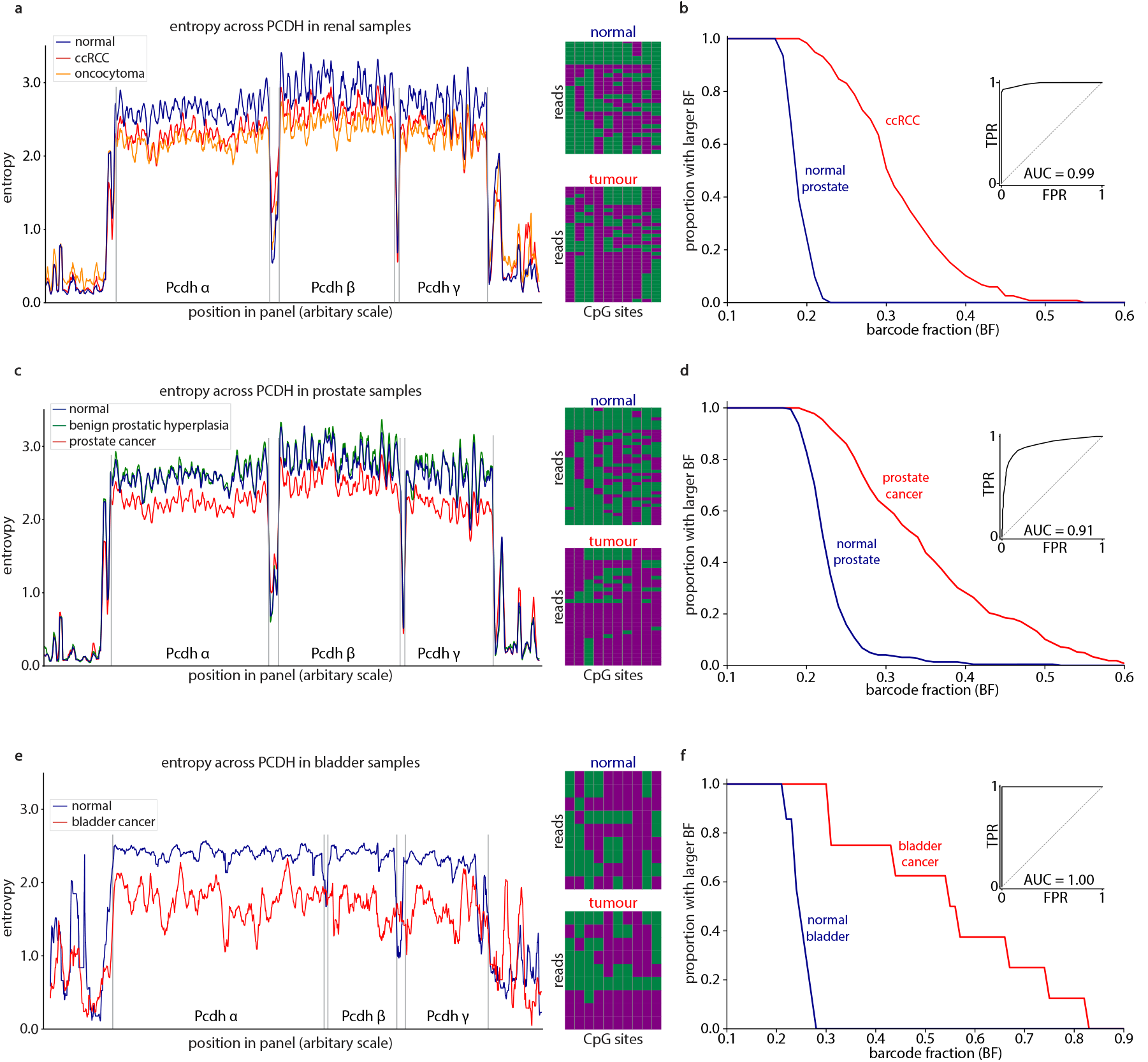
PCDH methylation patterns act as in-situ barcodes in other tissues. **a**. The drop in entropy in clear cell renal carcinoma samples (ccRCC, red) and oncocytomas (orange) compared to adjacent normal samples (blue). **b**. Reverse cumulative PCDH barcode fraction (BF) curves demonstrating that PCDH barcode fraction predicts ccRCC tumour status. **c**. The entropy across the protocadherin region in three sets of prostate samples, showing a reduction of entropy in cancer samples (red) relative to normal samples (blue) and non-clonal benign prostatic hyperplasia (green). **d**. Reverse cumulative PCDH barcode fraction (BF) curves demonstrating that PCDH barcode fraction predicts tumour status in prostate. **e**. The entropy across the protocadherin region in bladder, showing a reduction of entropy in cancer samples (red) relative to adjacent normal samples (blue). **f**. Reverse cumulative PCDH barcode fraction (BF) curves demonstrating that PCDH barcode fraction predicts tumour status in bladder.

Next, we investigated a dataset of prostate cancer biopsies (*n* = 267) and adjacent healthy tissue samples (*n* = 215) ^47^ and a small number of benign prostatic hyperplasias (BPH, n=12) expected to be polyclonal ^48,49^. The entropy in the prostate cancer samples was significantly lower than in the normal tissue samples (Mann-Whitney U test, *p<* 10^−6^), consistent with PCDH barcodes acting as clonal markers in prostate (Fig. 5d). Importantly, there was no drop in entropy observed in the hyperplastic samples, reflecting their polyclonality. As in renal samples, the average barcode fraction across the PCDH region can be used to classify tumours and normals with high accuracy (Figure 5f).

Finally, we considered a set of urothelial carcinomas (*n* = 8) and adjacent normals (*n* = 7) ^50^. In healthy bladder, PCDH had a high methylation entropy, which decreased in tumour samples (Fig. 5g). As in the other tissues, barcode fraction could be used to distinguish between healthy tissue and tumour with high accuracy (Fig. 5j).

Together, these observations demonstrate that methylation patterns in the PCDH gene cluster act as barcodes across a range of tissues and cell types and provide quantitative assessment of tissue clonality.

## Discussion

Here we have shown that the patterns of CpG methylation across the PCDH gene cluster act as a naturally evolving barcode that can be used for in vivo lineage tracing across multiple human tissues. In polyclonal samples, the PCDH region is characterised by a large diversity of methylation barcodes, which label large numbers of small clones. During a clonal expansion, the emergence of two clusters of barcodes inherited from the maternal and paternal haplotypes of the single founding cell results in a collapse in barcode diversity. This collapse provides a robust and quantitative measure of clonality that faithfully tracks the size and timing of expanding somatic clones.

The mechanisms that establish this remarkable diversity are not yet known as far as we are aware. Given the essential role of stochastic PCDH methylation in generating **the combinatorial diversity of isoforms required for neuronal** self-avoidance ^39–42^, it is tempting to speculate that patterns established early in development are inherited into both neuronal and non-neuronal tissues. While PCDH genes are not highly expressed in most non-neuronal tissues, suggesting the methylation states are functionally neutral, this requires further investigation. Understanding when and how this diversity is generated across different tissues is an important area for future work.

A key feature of the PCDH methylation barcode is its evolvability. We observed that the founding barcodes of a clone acquire further modifications through an ongoing process of epimutation, which we estimate to occur at a rate of approximately 0.1-3% per CpG per year. This constant generation of new diversity is a double-edged sword. On the one hand, it complicates lineage tracing, as the barcodes are not static. On the other hand, the evolvability is a useful feature, as it enables the resolution of subclonal events and the reconstruction of lineage dynamics within an established clone. The rate of diversification, which occurs over decades, is well suited to the timescales of somatic evolution in human ageing and cancer development. Further work is needed to assess how epimutation rates depend on acquired somatic variation.

PCDH barcodes currently have a number of limitations. The length of barcodes accessible via short-read sequencing (approximately 10 CpGs) restricts the theoretical diversity to only 1024 unique patterns, meaning the same barcode will independently arise multiple times within a large clone. This prevents a full phylogenetic reconstruction from a single time point. While we have shown that this low complexity can be overcome using long-read sequencing, our incomplete understanding of the epimutation process and higher rate of methylation sequencing errors currently make high-resolution phylogenetic reconstruction challenging.

Our findings demonstrate that PCDH methylation patterns act as in situ barcodes across multiple tissues, suggesting a pan-tissue phenomenon. We observed the same characteristic drop in methylation diversity in clonal tumours from kidney, prostate and bladder as we did in clonal expansions in blood. Data from lung suggests that methylation patterns in PCDH are similarly diverse there as in other tissues (supplementary note 19). This suggests that PCDH barcodes could find broad utility for assessing clonality in a wide range of contexts, from developmental biology to cancer. Because thousands of unique barcodes can be encoded in hundreds of bp of the PCDH locus, our work opens up the possibility of using low-cost PCDH amplicons to screen for individuals with clonal expansions, which are often the precursors to malignant disease. This could be useful in improving early detection of individuals at risk of developing future cancer.

## Methods

### Targeted duplex sequencing and enzymatic methylation sequencing data for UKCTOCs samples

This study used sequencing data generated for two previous studies by Watson et al. (Preprint 2024, targeted duplex sequencing) 51 and Fonseca et al. (Preprint 2025, targeted enzymatic methylation sequencing) 36. Briefly, both studies conducted deep targeted sequencing of serial blood samples collected from 50 women and age-matched controls annually up to 11 years prior to acute myeloid leukemia (AML) diagnosis as part of the UK Collaborative Trial of Ovarian Cancer Screening (UKCTOCS) (ISRCTN22488978; ClinicalTrials.gov NTC00058032) 45. For each blood draw (*n* = 530), somatic variants (SNVs and indels) were identified from a 58kb capture panel covering regions frequently mutated in haematopoietic malignancies to a mean duplexed depth of ∼ 1000X and used to infer clonal architectures in all donors. Targeted enzymatic methylation sequencing (EM-Seq) was also performed for each sample with a capture panel covering ∼ 200,000 CpGs at a mean deduplicated depth of ∼1000X. We used aligned bam files, methylation calls, and annotated variant calls from those studies in the methods described below.

### Germline variant calling, SNP phasing (paired-end alignments), and haplotype phasing (long-read alignments)

We used a custom workflow and scripts to SNP-phase paired-end targeted EM-Seq alignment data. Briefly, our workflow used Revelio software 52 to mask enzymatic conversions for downstream calling of germline single nucleotide polymorphisms. The resulting bam files were subject to standard SNP calling using GATK’s HaplotypeCaller tool GATK453 and the resulting vcf file was filtered to remove indels and homozygous SNPs. Next, we used the JVarkit tool, Biostar322664 54, to label read-pairs that overlap SNPs as either “ref” or “alt” in order to “SNP-phase” each read-pair. In cases where the read-pair covered multiple SNPs, we used the ref/alt status of only the first SNP covered by the read-pair to assign the SNP phase. We omitted putative heterozygous SNPs that were strand-associated - we hypothesize that these are artefacts from EM-seq conversion of homozygous G ⟶ A, A ⟶ G, T ⟶ C, or C ⟶ T variants.

To assign ONT sequencing long-reads to their respective haploid genomes, we used the Whatshap software package 55 integrated into the Clair3 workflow 56 using flow-cell appropriate ONT pre-trained models and the Longphase package (v1.7) 57. The resulting bam files include haplotype tags for each read that were used in the epiallele barcode extraction steps below.

### Construction of epiallele barcodes from paired-end EM-Seq alignments

For paired-end EM-Seq alignments from pre-AML donors, we used Bismark’s methylation extractor tool 58 to extract CpG methylation calls for each read pair in the bam alignment files. We developed a custom workflow to build SNP-phased epiallele barcodes for each read pair entry. Briefly, our workflow generated a dictionary structure in which each read pair was recorded in an independent entry that included vectors of CpG positions (genomic position) and states (0 or 1), as well as the read’s ref/alt status if it covered a previously called SNP (considering only heterozygous SNPs, and only the first if more than one was covered by a read pair). Of note, we considered each CpG as a single unit defined by the genomic position of the CpG on the positive (‘top’) strand, and thus converted CpG states and positions on reads with negative (‘bottom’) strand orientation to the corresponding positive genomic CpG position. In cases where the methylation state of a CpG was ambiguous (CpG dropout), the methylation state was assigned “nan” for the CpG position in the epiallele vector for that read.

### Discovery of high entropy loci

We defined the methylation entropy of a site to be

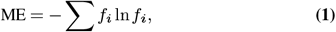

where *f*_*i*_ is the read fraction of a barcode with pattern i. For each site, we found the average entropy across all individuals where that site had a coverage of ≥100x. We excluded any sites that had sufficient coverage in fewer than half of the samples from subsequent analyses. Despite comprising only 2.8% of loci passing these filters, 10-mers in PCDH (hg19 chr5:140, 150, 000-140, 895, 000) accounted for 24% of the top 10% highest entropy sites.

### Quantification of methylation entropy across the panel

We considered the entropy across PCDH in all longitudinal blood samples which had passed sequencing quality control filters described in Watson et al. 38 and Fonseca et al. 36. For each unique barcode start position in the region, we found the mean 10-mer methylation entropy across all samples in which the site had a coverage of ≥100X. We then found the average entropy at each site in samples with and without large clonal sweeps (CF *>* 0.8 and CF *<* 0.2 respectively). To more clearly show the entropy pattern along the region, we plotted the rolling average over 30 adjacent epiallele start positions.

For the solid tissue datasets, the same analysis was performed, but with samples grouped according to tissue labels (‘benign’, ‘tumour’, etc.) instead of CF. All tumours were used, regardless of their purity. The lower sequencing depth for the kidney, prostate, and lung datasets necessitated using a lower minimum ten-mer coverage of 20X, while for the whole-genome bladder dataset a minimum depth of 10X was imposed.

### Filtering for high-confidence somatic variants

In addition to the filters described in Watson et. al. 51, we removed all variants that were present in a given individual at a VAF *>* 0.4 at all timepoints as being possibly germline. This approach removed variants which had previously been called as somatic from 12 individuals. Of these, four individuals were then called as having ‘missing drivers’ (C92_022, C92_023, C92_033, and C92_042) - we reasoned that these variants likely represented clones that had fully swept before sample collection began. Three had somatic variants which accounted for their BF (C92_034, C92_056, C92_062), and five had no evidence of a somatic clonal expansion from their methylation states or remaning genetic variants (C92_019, C92_061, C92_024, CNTRL_179, C92_068) - we reasoned that the largest variants in the latter two groups were likely germline. There were no expansions which were seen growing over time (and therefore certainly somatic) which were not marked by a corresponding increase in BF. We excluded all samples from CNTRL_192. This individual had a deletion which was covered by our panel for which the cell fraction could not accurately be measured. The cell-fraction against barcode-fraction relationship without the time-series germline filtering step is shown in extended data.

### Calculation of average Barcode Fraction

We defined the BF to be the read fraction of the two most frequent methylation patterns at a given locus. If the region did not experience ongoing epimutations during a clonal expansion, every cell in the clone would present the same two barcodes (one for each pattern), and therefore the BF would closely track the clonal cell fraction. We found the average BF across loci in the protocadherin region (hg19 chr5:140,150,020-140,895,000) excluding the constant region (hg19bbf chr5:140,300,000-140,400,000). Only loci that had a coverage of *>* 200x were included in the average, and only samples that had at least 100 (possibly overlapping) loci that passed these filterers were considered (the median number of loci included in the average of a sample was 1,370).

For the kidney, prostate, and lung datasets, where the depth was typically lower, the same procedure was followed but with a minimum depth of 20X, while for the low depth whole genome bladder dataset, a minimum depth of 10X was instead used.

### Simulation of expected cell-fraction barcode-size relationship

We characterised the relationship expected between clone sizes and BFs in in a large number (12,100) of sweeps simulated with realistic epimutation rates and random starting barcodes (see *Simulation of short-read barcodes during clonal expansions*). Simulations were run with a fitness *s* = 100%/year, a methylation rate drawn uniformly between 0/year and 0.1/year and demethylation rate drawn uniformly between 0/year and 0.01/year. The time between sweep initiation and measurement was drawn uniformly from 0 years and 40 years. To account for the non-uniform distribution of barcode patterns in non-clonal tissue, we quantified the background barcode-usage distribution in a sample with no sweep, and randomly selected reads from this distribution to represent molecules from wild-type cells in simulated individuals with incomplete sweeps. To simulate the effect of averaging across the region, ten ten-mers were simulated for each sweep, and the statistics averaged.

### Calling of samples with missing drivers

We called a sample as containing a missing driver if there was no simulated sweep (see *Simulation of expected cell-fraction barcode-size relationship*) within a cell fraction and barcode fraction of ±0.05. This procedure called 42 out of the 436 samples considered as having missing drivers, representing 16 out of the 99 individuals considered. Of these, nine samples, from four individuals, had variants which had been filtered out as possibly germline (*Filtering for high-confidence somatic variants*), leaving 33 samples, from 12 individuals without candidate drivers. To check these results were insensitive to the details of the epimutational process, we also investigated the effect of using a more stringent procedure to calling missing drivers, in which a sample is only called as having a missing driver if its BF *>* CCF and BF *>* 0.2. This metric quantifies which samples have BFs which are larger than either the somatic clone or background BF can account for, regardless of the epimutation rate. Of the 42 samples called as having a missing driver, 36 were called as having a missed expansion using this more stringent method. Of the 16 individuals who had a missing driver in at least one sample, only two (C92_012 and CNTRL_122) did not also have a missing driver called at some timepoint in using this stricter definition.

### Simulation of short-read barcodes during clonal expansions

We used a discrete-time simulation to investigate the effect of ongoing epimutations on the set of barcodes present during a clonal sweep. The simulation tracked the number of copies, *ns,p*, of a given pattern *p* present in cells with a given fitness *s* over time. Separately tracking barcodes with the same pattern but present in cells with different fitnesses allowed for complex dynamics, such a sub-clonal sweeps, to be simulated. Each time-step consisted of two stages, the first of which considered the effect of cellular dynamics. Cells were assumed to symmetrically divide at a constant rate *s* + 1*/τ* and die (or terminally differentiate) at rate 1*/τ*. In this model asymmetric divisions were ignored, as they have no impact on the set of barcodes present. The number of barcodes present in cells that symmetrically divide in the time period d*t* was therefore drawn from

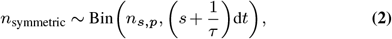

and number of barcodes present in cells that died was

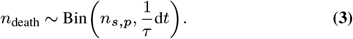

The updated barcode frequencies after this stage were then

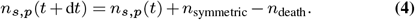

The second stage modelled the effect of epimutations on the barcode frequency. Barcodes were assumed to undergo methylating and demethylating epimutations at constant rates *µ* and *υ*. The number of cells in which a given unmethylated site became methylated was taken from

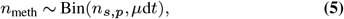

while the number of barcodes in which a given methylated site becomes unmethylated was sampled as

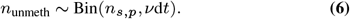

The number of copies of a given barcode was then updated according to

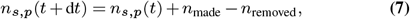

where *n*_made_ was the number of new copies of the barcode produced through epimutation from similar barcodes and *n*_removed_ was the number lost due to epimutations. Diploid cells were modelled by simulating a barcode of length 2*l* which was split into two barcodes of length *l* post-simulation. Simulation were initialised with the founding barcode pattern given a count 1 and continued for either a fixed time (for figure 2) or until the clone had fully swept (for inference of epimutation rates).

### Inference of epimutation rates

To measure the epimutation rates in clonal samples with a single timepoint, we developed a novel inference framework. We encoded the set of methylation patterns present at a given locus (a ‘pileup’) as a vector 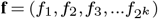, where *f*_*i*_ gave the read fraction of the barcode *i*. For each pair of rates considered (all pairs (*µ, υ*) where *µ* and υ*ε* took 128 values from 10*−*5 to 1, equally spaced in log space) we simulated a large number of pileups (min 506, max 624 for each pair) with randomly chosen starting barcodes. Our method quantified how closely an observed pileup could be reproduced in each of these simulated sets, using the Manhattan distance over read frequencies,

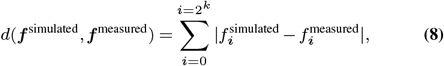

as the distance metric. To allow for high depth at every locus, and to be able to efficiently sample the space of starting barcode patterns, we used 6-mers for the inference. For each measured pileup, ***f*** ^measured^, we found the most-similar pileup, 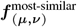 simulated with each pair of epimutation rates. We then quantified how consistent the measured pileup was with a pair of rates through

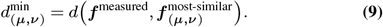

As any given pileup may be consistent with a range of epimutation rates, we found the average of 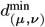 across the protocadherin region, excluding the constant region, for a given individual. We took the values (*µ, υ*) which minimised the average error to be the estimate of the rates. As pileups were simulated with a fitness of *s* = 100%/year, it was necessary to correct these estimates in sweeps with different fitnesses - theory and simulations show (supplementary notes 4B, 5) that the correct rescaling depends on the ratio of the assumed and true coalescence times for the sweep. In samples where the coalescence times are not known, the rates shown are calculated assuming a sweep time of 20 years, with possible ranges calculated based off the assumption that sweeps have a coalescence time of between 5 and 40 years. This method was applied on all clonal samples with a cell fraction *>* 80% without sub-clones observed above a 10% cell fraction. Simulations showed that this method can accurately reproduce ground-truth rates under such conditions - errors were calculated based off the uncertainty in the coalescence time and the uncertainty induced by applying this method to incomplete sweeps (supplementary note 5).

### Estimation of epimutation rates from individuals with sub-sweeps

For the two individuals (C92_023 and C92_033) who underwent sub-sweeps during the sampling period, the methylation and demethylation rates (*µ* and *ε* respectively) were calculated as

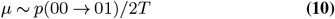

And

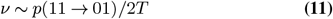

(supplementary note 4C), where *T* is the time elapsed between the start of the sweep and the start of the sub-sweep, and *p* is the proportion of sites undergoing a given transition. Sites were identified as having become methylated if they had a pre-subsweep average methylation of ≤0.3 and a post-subsweep methylation of ≥0.4, and having become demethylated if the corresponding quantities were ≥0.7 and ≤0.6 (changing these values made little difference to the estimated rate). While the initial coalescence times are not known for these samples, they are likely to be O(102) years. In C92_033, *p*_00→01_ = 15%, and *p*_11→01_ = 0.5%. This gives a methylation rate *µ* = O(10^−2^)/year and a demethylation rate ten-fold lower. For C92_023, the corresponding transition probabilities are 17% and 0.4%, giving the same order of magnitude epimutation rates.

### Long-read sequencing of myeloproliferative neoplasm patient and healthy donor samples

We acquired samples from Williams et al. (patient id: PD6466, peripheral blood granulocytes, myeloproliferative neoplasm, MPN) 33 and Mitchell et al. (donor id: KX004, peripheral blood mononuclear cells, healthy with early 15% clonal expansion) 32 and carried out whole genome, long-read methylation sequencing on the Oxford Nanopore Technologies (ONT) platform. We extracted 8-10 µg of DNA from cells using the QIAGEN DNeasy Blood & Tissue Kit and prepared sequencing libraries using ONT Ligation Sequencing Kits SQK-LSK110 (PD6646) or SQK-LSK114 (KX004) using manufacturer’s protocols. Sequencing was performed on PromethION 24/48 instruments using R9.4.1 flow cells to a mean depth of ∼282.7X (PD6646) or R10.4.1 flow cells to ∼150X (KX004). Base calling (including 5mCG modifications) and alignment from pod5 files was done using the ONT dorado workflow (ONT-models: dna_r10.4.1_e8.2_400bps_sup@v4.3.0 and dna_r9.4.1_e8_sup@v3.3.0, for KX004 and PD6646, respectively) using the GRCh38 reference build. The resulting bam alignment files, which included CpG modification calls, were subjected to germline variant calling and haplotype phasing as described below.

### Construction of epiallele barcodes and barcode matrices from haploid-phased long-read sequencing alignments

We used the ONT Modkit tool (“Modkit extract calls” with default settings) to extract CpG methylation calls for each read in the ONT long-read bam alignment files.

We developed a custom workflow to build haplotype-phased epialleles for each read entry in the bam files. Briefly, our workflow generated a dictionary structure in which each read was recorded and an independent entry that included vectors of CpG positions (genomic position) and states (0 or 1) and the haplotype assigned to that read by Whatshap. In cases where the methylation state of a CpG was ambiguous (CpG dropout), that CpG position was omitted from the epiallele vector for that read.

Next, using custom scripts we constructed barcode matrices by sampling long-read barcodes across the PCDH gene cluster in ∼3-exon windows. Due to the large fraction of CpG dropout in long-read sequencing (20-30%), constructing a submatrix of reads (rows) and CpGs (columns) from a pileup of each window in which all selected CpGs are called poses a non-trivial, NP-Hard computation challenge (e.g. for 500 reads covering 500 CpGs a brute force approach over all subsets would require 2500 possible configurations). We therefore implemented a greedy approach where matrices were constructed using an iterative process to maximize the number of covered CpGs while retaining an informative number of barcodes (i.e., target of >50-100 reads). We required ≤20% CpG dropout and ≤5 contiguous dropouts for contributing reads with a target of >50 CpGs per matrix. Through trial and error we found that semi-sliding windows covering an average of 29.5kb and 646 CpGs each was required to meet these matrix dimensional requirements (see Supplementary Table S1). Although the average read length in PD6646 and KX004 was 10.3kb and 3.8kb, respectively, which covered an average of 220 and 81 CpGs per read in the PCDH region, the final haploid matrix size was significantly reduced due to CpG dropout. For PD6646 (266X mean sequencing coverage across PCDH), this approach produced matrices with means of 66.1 (range: 38-98) barcodes and 152.2 (range: 29-280) CpGs per haploid. For KX004, which had 121X mean sequencing coverage (target of >25-50 reads per matrix), matrices had means of 34.2 (range: 15-76) barcodes and 380.0 (23-380) CpGs per haploid genome.

### Clustering of barcode matrices and calculation of cluster read fractions

Long-read barcode matrices subjected to hierarchical clustering based on the premise that, for each haploid matrix, (i) the sizes of clusters formed on the basis of barcode similarity would reflect the sizes of clonal subfractions, (ii), the hierarchical similarity of the barcodes would recapitulate phylogenetic relationships between (sub)clones, and (iii) one haploid matrix would serve as a cross-validation to the other haploid matrix in the same genomic region (PCDH window). In order to carry out hierarchical clustering on each matrix given CpG dropout, it was necessary to impute missing CpG calls using a k-nearest neighbours (kNN) approach based on Hamming distance between barcodes (k=3). For each matrix, this method compares each barcode (row) to all others using Hamming distance which is computed using only CpG positions (column) where both barcodes have non-’N’ values. Next, the k most similar rows are selected (lowest normalized Hamming distance) and the missing value is determined using the majority vote across those k rows at that column, with fallback to the column-wise majority when neighbour information was unavailable. We first assessed the effect of imputation on matrix clustering by comparing complete matrices generated by an agent-based cell growth simulator (see section below *Simulation of longread CpG barcodes in a growing stem cell population* and Supplementary Note B) to the same matrices with varying amounts of random CpG dropout (see Supplementary Fig. S8). We determined that imputation using Hamming kNN with 20% non-adjacent dropout was not detrimental for accurate downstream barcode clustering (∼80% and ∼65% imputation accuracy for clonal cell fractions of 80% and 20%, respectively, see Supplementary Fig. S8a). We applied the Hamming kNN method to impute missing CpG states in the haploid barcode matrices generated from long-read data (having <20% non-adjacent CpG dropout) as described above. For each of the resulting matrices, we carried out hierarchical clustering using Hamming distance and average linkage (UPGMA) without additional scaling or normalization, and calculated the fraction of reads assigned to (i) the two clusters resulting top-level bifurcation, and (ii) the with a five cluster cut-off. We reasoned that a monoclonal expansion starting from a random wild-type barcode pool with 1% epimutation rate would allow discrimination of the healthy and cancer fractions at the first branch. We further reasoned that given the number of barcodes (50-100 per haploid) we would not be sufficiently powered to discriminate beyond ∼5 clades.

### Simulation of long-read CpG barcodes in a growing stem cell population

We constructed an agent-based simulation framework to model clonal expansion (Wright–Fisher process) and long-read epiallele barcodes in a fixed-size stem cell population over time. The simulator tracks individual stem cells and their epigenetic states (haploid epiallele barcodes) while incorporating stochastic cell selection, clonal mutations, subclonal fitness advantages, and epimutational noise. The population was initialized with *N* = 10000 wild-type stem cells, each represented as a digital agent. At each simulated time step (defined as one year), the entire population was replaced by sampling *N* offspring from the previous generation using fitness-proportional selection with replacement (Moran-like process), thereby maintaining homeostatic population size. At selection initiation, a single randomly selected cell was assigned a founder clonal mutation with a fitness advantage *s*_1_ (probability of producing offspring to (1 + *s*_1_)). After time (*T*) a subclonal mutation with additional advantage *s*_2_ was introduced in a random descendant of the founder clone. The cumulative fitness for subclonal descendants was therefore (1 + *s*_1_ + *s*_2_). Each cell possessed a diploid epigenetic state represented by two haplotypes, each with 500 independent CpGs comprising binary methylation values (0 = unmethylated, 1 = methylated). These haplotypes were initialized randomly under a binomial model (p = 0.5). During cell division, epialleles were inherited from the parent with the potential for stochastic epimutations applied to each CpG site: 0 − 1 with probability *µ*, and 1 − 0 with probability *ϑ* (default: *µ* = *γ* = 0.01). These epimutational events were applied independently to each CpG site per haplotype per generation. For each simulation year, the full population’s diploid epialleles were recorded and stored. Haploid selection was performed randomly per allele and stored alongside their original haplotype origin (“A” or “B”). Clonal and subclonal fractions were quantified at each time step based on the cumulative proportion of cells carrying the founder or subclonal mutation.

To assess the impact of barcode length on clonal reconstruction, as well as the accuracy of Hamming kNN imputation of missing CpG states due to sequencing dropout, we first subsampled the barcode population at ∼70% founder (∼60% subclone) clonal cell fraction by randomly selecting 100 barcodes belonging to a single haploid (e.g., all haploid “A”,Supplementary Fig. Aa). We then introduced random methylation calling errors (10% of sites, “noise”) and random call dropout (for 20% of calls, we replaced methylation state, 0 or 1, with “nan”). Next, we titrated barcode length by selected the first 10, 25, 50, 75, 100, or 200 CpGs in each barcode. We then carried out Hamming kNN imputation (as described above) on each matrix and subjected the resulting barcodes to hierarchical clustering. We visually compared the topography of the ground truth clustering (Supplementary Fig. Ab) to the clustering generated for matrices at length (Supplementary Fig. Ac).

We further tested the impact of dropout imputation by constructing matrices from randomly subsampled haploid “A” barcodes (*n* = 100) with 100 CpGs each and 10% random noise at low (∼20%) and high (∼70%) mutant founder clonal cell fractions (Supplementary Fig. Bb,c, top “ground truth trees”). We simulated dropout rates ranging from 1% to 50% for all matrices and assessed imputation accuracy (percent of correctly imputed CpG states) by comparing imputed CpG states to pre-dropout states (Supplementary Fig. Ba). Note that at high clonal cell fraction (i.e., ∼70%) missing values could be reliably recovered from neighbouring barcodes that share recent ancestry, leading to robust assignment of cells to the correct clonal family. In contrast, at smaller clonal cell fraction (i.e., ∼20%), imputation accuracy dropped to 60-65%, consistent with reduced redundancy and fewer closely related neighbours available for inference. This suggests that imputation is most reliable for preserving correct barcode-to-clone assignment in dominant clones, whereas clonal assignment becomes progressively less accurate for smaller expansions or subclones, particularly under high dropout. To test this, we hierarchically clustered these matrices at dropout rates of 0%, 5%, 10%, 20%, and 40% and compared topology and accuracy of clade/cluster assignment to the ground truth trees (Supplementary Fig. Bb,c). Despite reduced imputation accuracy at the most extreme parameter values tested (20% cell fraction and 40% dropout), barcodes were correctly assigned to their primary clonal branch.

### Identification of loci with most-ideal behavior

All short-read numerical results are based off averages over all loci with sufficient depth outside of the constant region. For illustrative purposes (eg Supplementary Figs. S16, S17), it was occasionally useful to restrict to a set of most-optimally behaved loci. To discover such a set, we found the cell-fraction top-two statistic relationship for each elocus in the region that had at least 100 samples with a depth of at least 100. We then computed Spearman’s rank correlation coefficient for each of these eloci. This measured how monotonic the relationship between the clone size and barcode size is. We took the 119 loci (4.9% of total) with the largest Spearman’s rank to be the most-optimal set of sites.

## Acknowledgements

We thank E.Joanna Baxter, Emily Mitchell and Jyoti Nangalia for sharing material from PD6646 and KX004 samples. We thank Tom Maniatis and his lab for useful input on PCDH biology. We thank Calum Gabbutt and Heather Grant for advice on tree-building from CpGs. We thank Florian Merkle, Daniel Fisher, Alex Cagan, Ugne Vaitkute, Barbara Walkowiak, Alex Frankell, and all members of the Blundell lab for helpful comments. We are indebted to the generosity of the women who participated in UKC-TOCS and made their samples and data available for secondary research.

J.R.B, C. T. B and S.F.H are supported by the Early Cancer Institute, the CRUK Cambridge Centre and the NIHR Biomedical Research Centre. J.R.B. is supported by a UKRI Future Leaders Fellowship (MR/S031782/1). S.F.H is supported by an International Alliance for Cancer Early Detection (ACED) PhD studentship. A.V.A.F and C.B are both supported by ACED pathway awards from the International Alliance for Cancer Early Detection. C.J.W. was supported by a CRUK Clinical Research Fellowship and is currently supported by a Wellcome Early Career Award (226929/Z/23/Z). U.M., S.A. and A.GM. are supported by the NIHR UCL Hospitals Biomedical Research Centre and by Medical Research Council Clinical Trials Unit at UCL core funding (MR_UU_12023).

## Author Contributions

J.R.B conceived of the study. S.F.H and C.T.B performed the relevant analyses with input from J.R.B. A.V.A.F, C.J.W, A.T generated data used in this study. A.D.R.Y. provided early evidence of PCDH as a barcode. H.M.L.B. performed the single-timepoint analysis of pre-AML with input from S.F.H, C.T.B and J.R.B. H.D and H.L.Y provided data from prostate. U.M, S.A and A.G.M provided material and advice on study design. The manuscript was written by J.R.B, S.F.H and C.T.B with input from all authors.

