## Supplementary materials for "In vivo lineage tracing across human tissues using methylation barcodes in the protocadherin gene cluster"

### Supplementary Note 1: Construction of epiallele barcodes from paired-end EM-Seq reads

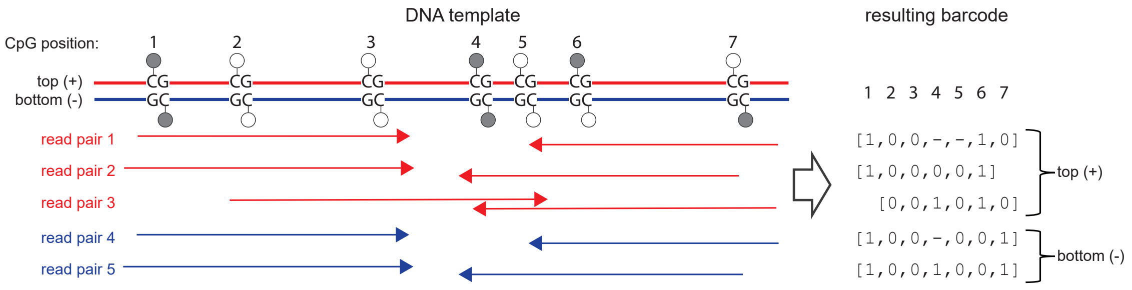


**Fig. S1. Schematic of encoding epiallele barcodes from paired-end EM-Seq CpG methylation calls**

### Supplementary Note 2: Ruling out alternative explanations for PCDH methylation behaviour

**Cell type effects.** To establish that PCDH methylation barcodes act as in-vivo lineage markers, it is important to rule out the possibility that cell-type effects give rise to the phenomena we observe. Numerous observations show that such an interpretation is not consistent with the behaviour of methylation patterns in PCDH. Most importantly, as reported in Fonseca et al.[36], the majority of the DNA sequenced from UKCTOCS peripheral blood samples comes from Neutrophils, and the cell-type changes between samples with and without large clones are too subtle to account for the dramatic drop in diversity observed during clonal sweeps. Furthermore, the barcodes that expand at a given locus are chosen randomly (see supplementary note 14), whereas if barcodes marked cell type, one may expect the same methylation patterns to be found at a given site across individuals. The tri-modal methylation distribution function post-sweep is also not consistent with cell-type effects - there is no reason that cell-type biases would cause a given CpG locus to switch to being fully methylated, fully unmethylated, or methylated in half of reads. Finally, the behaviour of the barcodes during a sub-clonal event (in which a new, slightly altered set of barcodes establish) would not be expected if the region was strongly cell-type associated.

### Supplementary Note 3: Methylation distribution of barcode CpGs in the PCDH gene cluster

Previous work by our group and others found that the methylation distribution of fluctuating CpGs (fCpGs) reflects clonal diversity of blood stem cells where a “W"-shaped distribution emerges as a clone expands[34–36]. Methylation distributions of PCDH barcode CpGs closely matched those that we previously observed in an independent set of fCpGs for these samples and distribution variance was similarly quantitative for clonal cell fraction in cases with detectable drivers (Supplementary Fig. S2a,c,d). The clonality statistic for PCDH barcodes also closely correlated with the distribution variance of non-PCDH fCpGs (Supplementary Fig. S2e). We also observed similar differences in homogeneity of CpG methylation states in expanded versus non-expanded donors, which is consistent with previous findings cited above (Supplementary Fig. S2f). Specifically, methylation state homogeneity is high in diverse, non-expanded cell populations and low in oligoclonal and expanded populations. Supplementary Fig. S2f shows homogeneous methylation regions in non-expanded samples as mostly white space in contrast to the more heterogeneous behaviour of samples with high clonal cell fractions. These patterns also highlight donors with predicted missing drivers, the tendency of epimutations toward methylation, and potential CpG positions that may be less mutable across the region. Positions that tend to be consistently (un)methylated across all donors are candidates for further filtering in future studies.

The behaviour of CpGs used in our PCDH barcodes closely resembles that of previously identified fCpGs in other regions of the genome in a similarly quantitative manner, further supporting their utility to detect clonal expansions and as clonal lineage markers. Although it is unclear whether similar mechanisms drive CpG methylation in the PCDH gene cluster, they nevertheless appear to have similar properties as they synchronize during clonal expansion. Furthermore, a region of high-density CpGs with this behaviour allows for both less-expensive, deeper targeted sequencing and the benefit of the additional information from phasing.

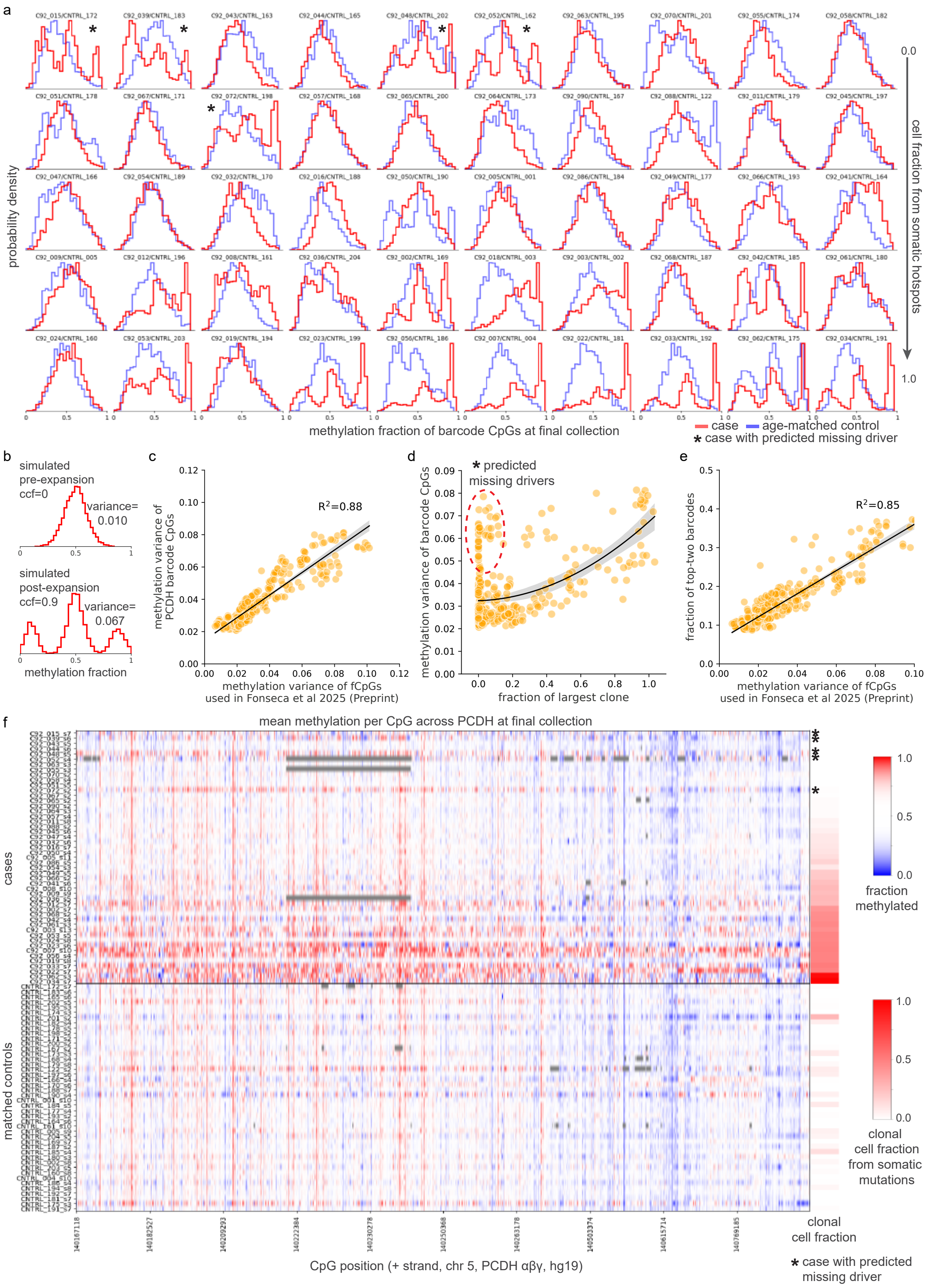


**Fig. S2. a.** Distribution of CpG methylation states in barcode CpGs from the PCDH gene cluster region at the last time point collected sorted by increasing clonal cell fraction (CCF) as determined by somatic mutations (case=red, control=blue). **b.** The distinct "W"-shaped distribution that arises from synchronization of CpG states during clonal expansion is observed in PCDH CpGs, the variance of which correlates well with previous work[36] using an independent set of fluctuating CpGs (fCpGs) in the same sample set (**c**). This variance quantitatively reflects the cell fraction of largest clone (as determined by somatic mutations) (**d**) and is highly correlated with the top-two barcode fraction (BF) (**e**). **f.** Methylation patterns across the PCDH region are shown for cases (top, sorted by detectable mutation CCF) and matched controls (same order, bottom) and are consistent with other studies using array-based fCpGs measurements[35] where non-expanded stem cell populations exhibit homogeneous states (i.e., average ∼0.5, white) and expanded and oligoclonal population state are heterogeneous; either methylated (1, red) or unmethylated (0, blue). Cases with predicted missing drivers (asterisks) have patterns similar to those with detectable higher frequency somatic mutations (red clonal cell fractions)

### Supplementary Note 4: Modelling barcode dynamics

**A. Evolution of the set of barcode fractions during a clonal sweep.** We consider the evolution of the cell fraction $f$ of the barcode patterns present during a somatic sweep. New copies of a given barcode can arise through either symmetric cell divisions or epimutations from closely related barcodes, while existing copies can be lost due to epimutations. For a given methylation rate $\mu$ and demethylation rate $\nu$, the dynamics of the barcode fractions $f_b$ are therefore given by

$$\frac{df_b}{dt} = S(t)f - (\mu n + \nu(l-n))f_i + \mu \sum_{i\in D_b} f_i + \nu \sum_{i\in M_b} f_i, \quad \textbf{(12)}$$

where $M_b$ and $D_b$ are the sets of barcodes which are a single methylation or demethylation edit away from barcode b, and $S(t)$ is the instantaneous growth rate of the clone. When barcodes are sufficiently long, any individual pattern will be rare outside of a clonal expansion. The barcode fraction within a clone, $\tilde{f}$, will then be given to a close approximation by $\tilde{f} \approx f/\mathrm{CF}$, where $\mathrm{CF} = \mathrm{CF}_0 \exp(\int S(t')dt')$ is the clonal cell fraction. It then follows that

$$\frac{d\tilde{f}_b}{dt} = \tilde{f} - (\mu n + \nu(l-n))\tilde{f}_i + \mu \sum_{i\in D_b} \tilde{f}_i + \nu \sum_{i\in M_b} \tilde{f}_i. \quad \textbf{(13)}$$

The set of barcode fractions expected, $\tilde{\boldsymbol{f}} = (\tilde{f}_1, \tilde{f}_2, ...)$, is then governed by

$$\frac{d\tilde{\boldsymbol{f}}}{dt} = \mathrm{M}\tilde{\boldsymbol{f}}, \quad \textbf{(14)}$$

where the elements of M are defined in equation 12, $n$ is the number of unmethylated sites in the barcode, and $l$ is the barcode length. The above can be solved using a matrix exponential,

$$\tilde{\boldsymbol{f}}(t) = \exp(\mathrm{M}t)\tilde{\boldsymbol{f}}_0. \quad \textbf{(15)}$$

Hence, the predicted barcode dynamics within a clone depend on epimutation rates and coalescence time only through the products $\mu t$ and $\nu t$ and is independent of the clonal cell fraction $F(t)$. This allows for inference of epimutation rates even when $F(t)$ is non-exponential, for example due to clonal competition. While previous studies[35] have claimed to find independent estimates of $\mu$, $\nu$, and $t$, such work is based on strong assumptions about the state of fCpG sites at birth which are difficult to verify with existing data. To infer epimutation rates it is therefore necessary to use pre-existing estimates for the initiation time based on clonal growth rates.

**B. Read fraction of founding barcode during a clonal sweep.** In the limit of low $\mu t$, $\nu t$ the effect of epimutations back to the founding barcode will be negligible and the frequency of the founding barcode will be given to a close approximation by

$$\frac{d\tilde{f}}{dt} = -(\mu n + \nu(l-n))\tilde{f}. \quad \textbf{(16)}$$

We can solve the above to get

$$\tilde{f} = \exp(-(\mu n + (l-n)\nu)t), \quad \textbf{(17)}$$

where $\tilde{f}(t=0) = 1$. Therefore

$$f(t) = \mathrm{CF}(t)\tilde{f}(t) = \mathrm{CF}(t)\exp(-(\mu n + \nu(l-n))t). \quad \textbf{(18)}$$

Again, the barcode fraction within a clone depends only on the products $\mu t$ and $\nu t$.

**C. Effect of ongoing epimutations on longitudinal barcode frequency plots.** The following plots illustrate the barcode behaviour expected during a clonal sweep and sub-sweep. If there were no ongoing epimutations, the two founding barcodes would completely sweep with the clone (Supplementary Fig. S3a). For epimutations at a rate $\mathrm{O}(10^{-2})\mathrm{year}^{-1}$, founding barcodes continually epimutate into barcodes with similar patterns, and so after a sweep two families of closely related patterns establish. When there is a bias in the epimutation rate, less methylated barcodes generally reach lower read fractions, as unmethylated sites undergo more epimutations than methylated sites (Supplementary Fig. S3 c). The same behaviour can be observed in clonally expanded UKCTOCS blood samples (Fig. 3 c,d,e, Supplementary Fig. 14a). The founding barcode on a haploid for a sub-sweep may or may not have undergone an epimutation relative to the founding barcode of the initial clonal expansion. The subsweep can therefore be tracked by both, only one, or by neither haploid (Supplementary Fig. S3 d,e,f respectively), depending which of the two founding barcodes of the sub-sweep have undergone epimutations relative to the founding barcodes of the original sweep. All three behaviours are observed in samples from UKCTOCS blood donors with subclonal expansions during sample collection (see Supplementary Fig. S17 14016710,140167242 for detection by both haploids, 140167805, 140209211 for detection by only one haploid, and 140182410, 140167560 for no detection).

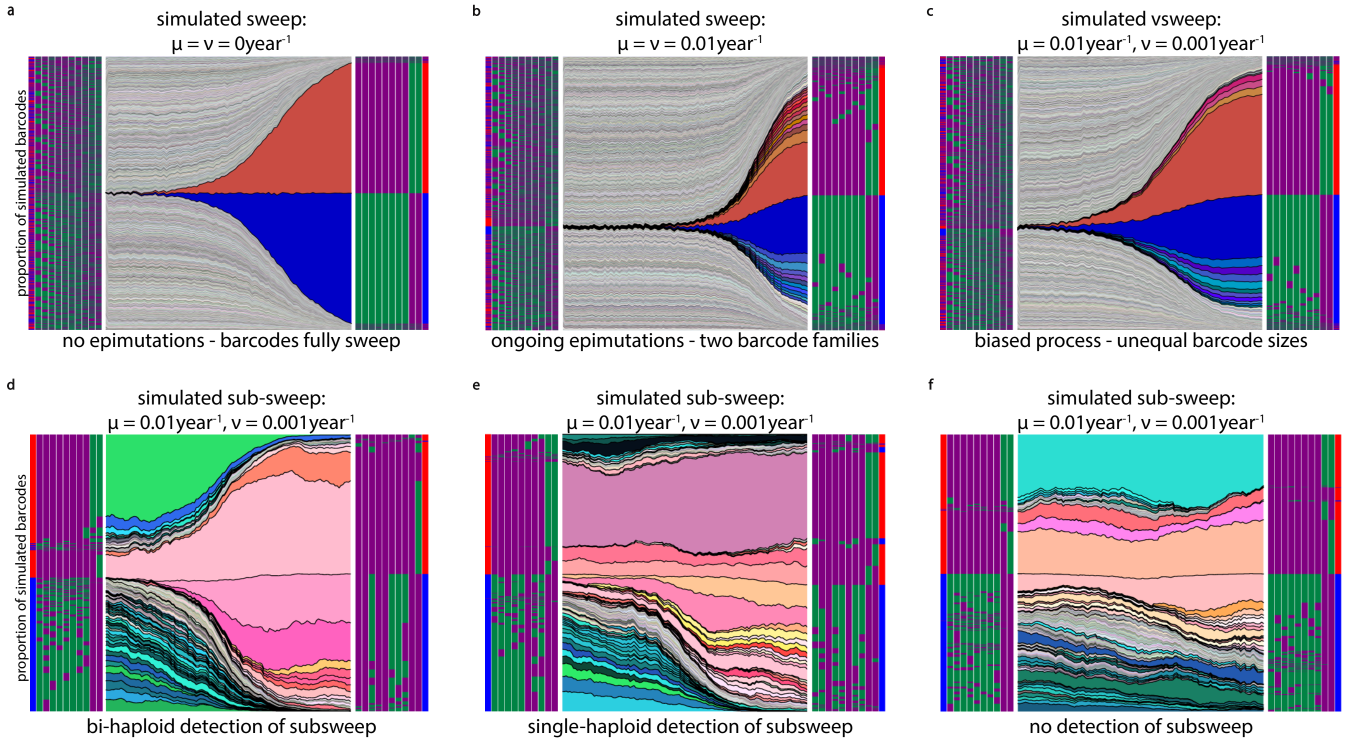


**Fig. S3. Effect of ongoing epimutations on barcode behaviour during sweeps and sub-sweeps. a.** A simulation showing the growth of the two founding barcodes during a clonal sweep with no ongoing epimutations. **b.** The same with ongoing methylating and demethylating epimutations at a rate of $0.01/$year. textbfc. A simulation showing the growth of barcodes under a realistic epimutation process. **d,e,f.** Simulations run with the same starting barcodes, sweep fitnesses, and times but different random seeds, showing three possible behaviours of barcodes during a sub-sweep - detection by both haploids, just one haploid, or no detection respectively.

### Supplementary Note 5: Benchmarking inference on simulated data

To investigate the robustness of our inference framework to realistic complications, we benchmarked its ability to recapitulate ground-truth epimutation rates in a range of simulated scenarios. To test if we could accurately measure the epimutation rate in expansions with different fitnesses and wait times, we considered a simulated cohort of clonal sweeps with a range of fitness drawn uniformly from $0.5 - 2/$year and sampling times from clone initiation drawn uniformly from $5 - 15$years. We only considered samples with a simulated CF $\geq 0.9$. Consistent with theory (supplementary note 4), we found that we could accurately determine the epimutation rate times the coalescence time under such conditions. This allows us to infer the epimutation rates when the coalescence time is known, and bound the rates when the time is not known.

To investigate the effect of incomplete sweeps on the estimated rates, we considered the sweeps which had $0.8 \leq$ CF $< 0.9$. We found that, because at small epimutation rates the residual diversity will be dominated by the remaining wild-type barcodes, this leads to an overestimate if the rates are small. We therefore only applied our method to UKCTOCS blood samples with sweeps with CF $> 0.8$, and estimated the uncertainty caused by incomplete sweep sizes by finding the range of ground-truth rates that gave such a measured value in-silico.

Finally, to investigate the effect of sub-clonal expansions on the measured epimutation rates, we simulated the set of patterns produced by a sub-clonal growth growing on the background of a full sweep. When large sub-clones were present (sub-CF $> 20\%$), the obtained epimutation rates were underestimates of the ground-truth rate (supplementary Fig. S4c). We reasoned that such behaviour arises because the barcodes within the sub-clone have had less time to epimutate, giving less residual diversity than would be expected, giving an underestimate of the rates. We therefore limited our analysis to individuals with sub-clones of a maximum cell fraction of $10\%$. Having sub-clones smaller than $10\%$ had a negligible impact on our ability to accurately infer rates (supplementary Fig. S4d). We did, however, run inference on samples which had sub-clones of a similar size to the main clone (for example, as present in C92_002_s7). For these samples we used the coalescence time of the sub-clone.

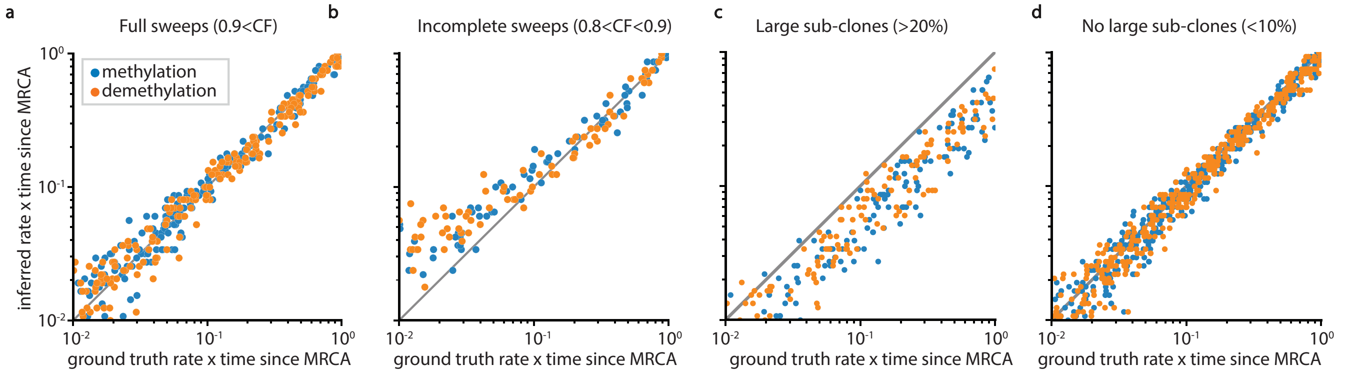


**Fig. S4. Benchmarking of epimutation rate inference**. **a.** Example inference for sweeps simulated with a range of fitnesses and wait times, with cell fractions $> 90\%$. **b.** Inference under the same conditions, but with cell fractions between $80\%$ and $90\%$. **c.** Impact of large sub-clones (sub-CF$> 20\%$ on inferred rates. **d.** Impact of small sub-clones (sub-CF$< 10\%$) on inferred epimutation rates.

### Supplementary Note 6: PCDH methylation barcodes reflect sweep timings

Under our model of barcode dynamics, the read fraction of the largest barcode within a clone at a given locus decreases in time as $\exp(-(\mu n + \nu(l-n))t$. Under the assumption that the epimutation rates $\mu$ and $\nu$ are constant over time, methylation barcodes should therefore encode information about the coalescence time for a sweep. We found that the BF was larger in samples C92_023 and C92_033 after a sub-sweep than before it, suggesting that the coalescence time for the parent sweep was longer than that for the sub-sweep, consistent with the general pattern of fitness effects observed in clonal sweeps with increasingly many driver mutations[(38)]. To ensure that this observation was not confounded by biases in the barcodes of origin for the sub-sweeps, we found the mean read fraction of each possible 6-mer pattern conditioned on it being one of the largest two at a given locus. The same patterns generally had higher read fractions post sub-sweep that pre sub-sweep.

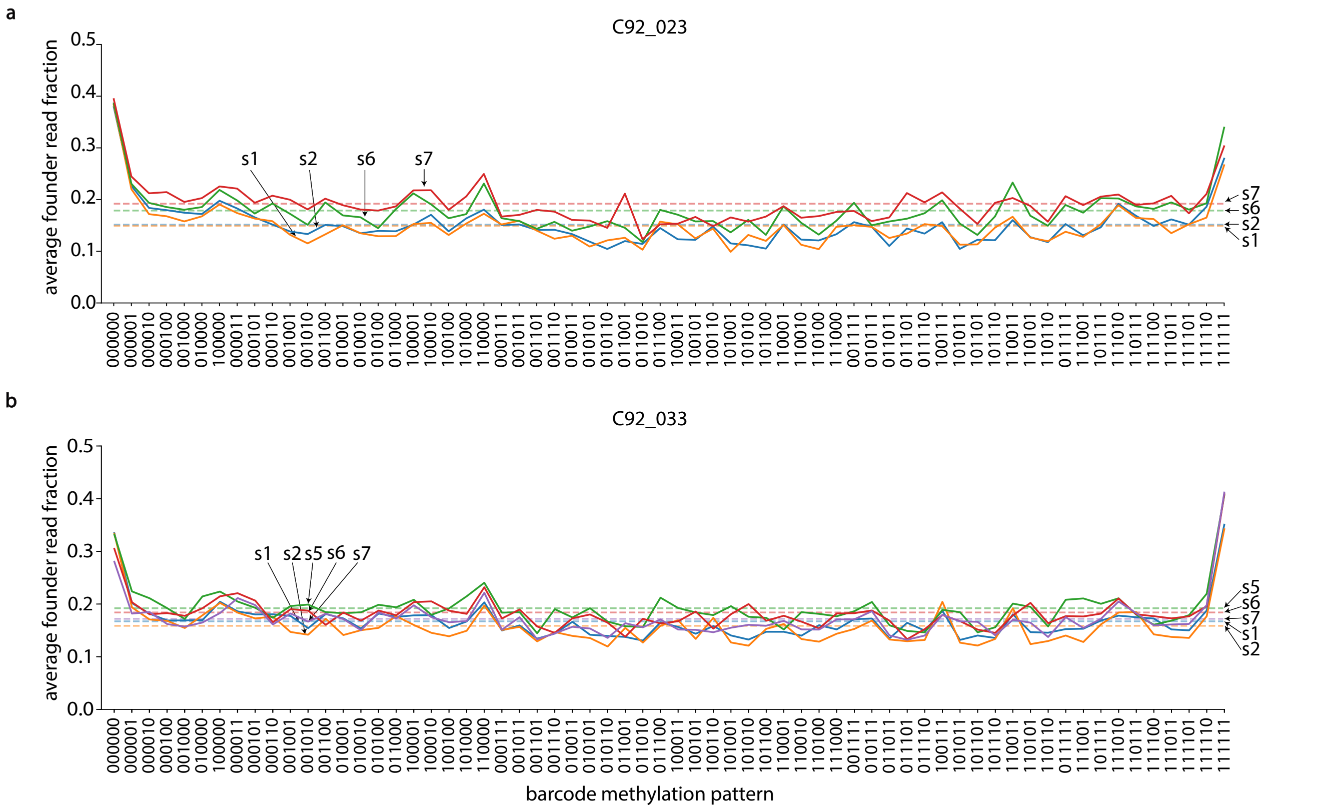


**Fig. S5. PCDH methylation barcodes reflect sweep timings. a.** Average read fraction of largest barcodes against the pattern of the largest barcode for C92_023. **b.** The same for C92_033.

### Supplementary Note 7: Other methylation barcode regions

We found that an extended region of chromosome 16 (hg19 chr16:32214053-34208607) had a lower methylation entropy in individuals with large clonal sweeps than in polyclonal samples (supplementary Fig. S6a, red and blue lines respectively). While sites in this region have high methylation entropy, they are less diverse than those in PCDH (S6b). Unlike in PCDH, we are not aware of any underlying reason for this high methylation diversity. Nevertheless, the average BF across this region tracks ground-truth cell fractions (supplementary Fig. S6c), and is strongly correlated with the same quantity in PCDH (supplementary Fig. S6d). Controls showed more diverse patterns of methylation than cases (supplementary Fig S6e,f), although the set of patterns in polyclonal samples was typically less diverse than in PCDH. The dynamics of methylation patterns at single loci in this region bore some resemblance to the dynamics of somatic variants over time. We found that methylation barcodes in this section of chr16 typically could not be used to track sub-clonal events - often the barcodes in these regions had become almost fully saturated as fully methylated (or in some cases fully unmethylated), limiting their utility for high resolution lineage tracing. Nonetheless, the existence of another extended region in our panel which displays some of the same features as PCDH raises the possibility that there may be many such regions across the human genome.

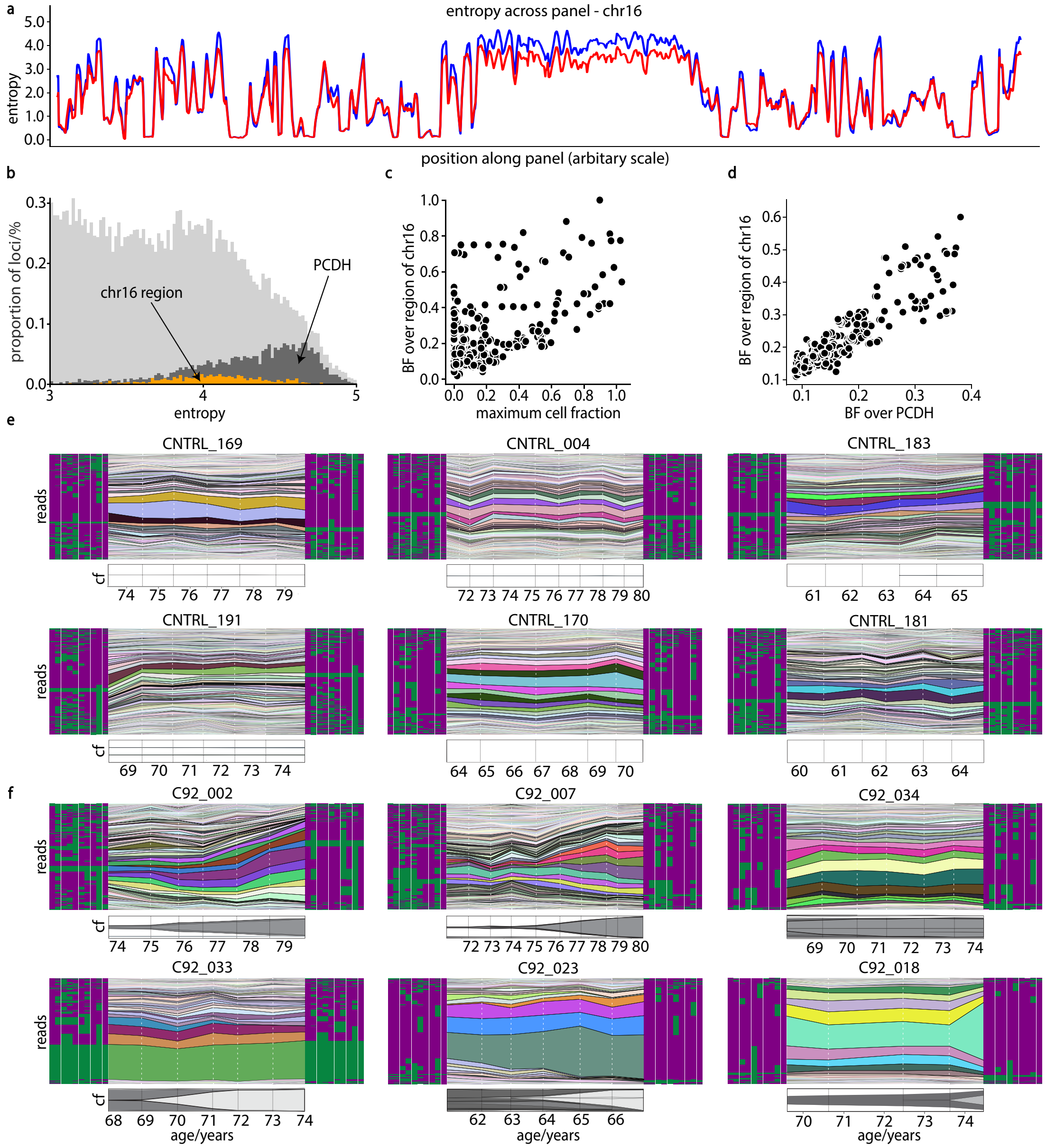


**Fig. S6. A region of chromosome 16 acts in a similar way to PCDH a.** The 10-mer methylation entropy of individuals with (red) and without (blue) large clonal sweeps in the targeted EM-seq panel across chromosome 16. **b.** The top-end of the 10-mer methylation entropy distribution, with PCDH and the region of chr16 highlighted. **c.** BF computed across the region of chr16 against the maximum cell fraction found in somatic sequecing. **d.** The average BF over the region of chr16 against the average BF in PCDH. **e.** Barcode frequencies over time in controls with no observed somatic sweeps. **f.** The same for cases with high-frequency somatic variants.

### Supplementary Note 8: Simulation of long-read stochastic CpG barcodes in a growing stem cell population

#### A. Performance of simulated long-read barcode matrices.

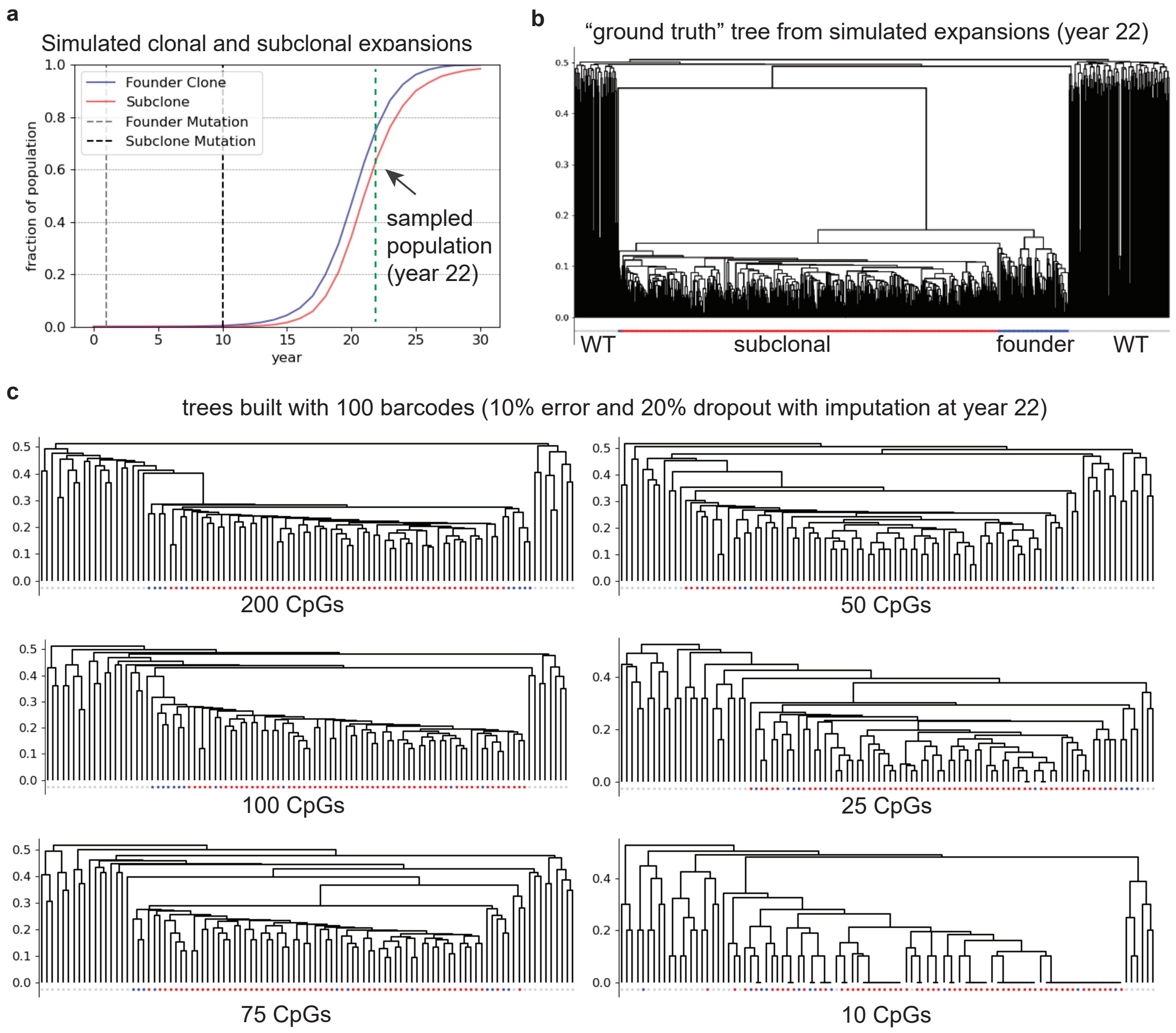


**Fig. S7. Impact of barcode length on clonal reconstruction. a.** Simulated stem cell population with founding (year=1) and subclonal (year=10) mutation events (see methods, Simulation of long-read CpG barcodes in a growing stem cell population; N=10k diploid cells, initial barcode states were random, epimutation rate=$0.01$/year). The population was sampled 22 years after initiation of the mutant clone (s=0.5), which allowed 12 years of subclonal expansion as well (s=0.5), at $\sim$70% CCF (subclone $\sim$60%). **b.** Ground truth tree showing wild-type (WT) and linear clonal (founder, blue) and subclonal (red) cells. **c.** Trees constructed using 100 barcodes (haploid "A" only) with various numbers of CpGs per barcode. Note, after subsampling barcodes from the population, they were subjected to the addition of 10% random error (noise) and 20% random dropout followed by imputation for hierarchical clustering (see methods).

### B. CpG dropout imputation accuracy and its affect on cluster assignment.

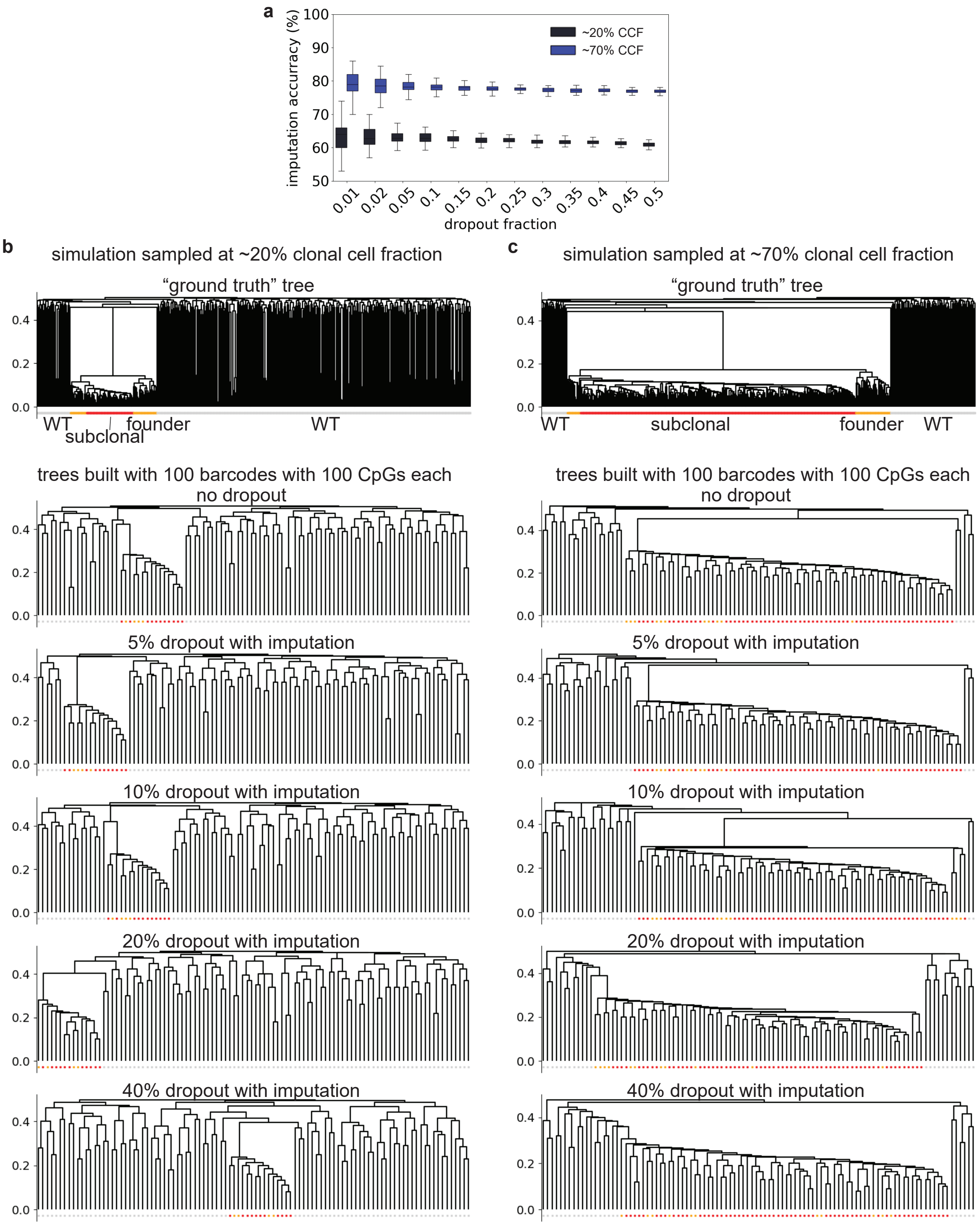


**Fig. S8. Impact of CpG dropout and Hamming kNN imputation on clonal reconstruction. a.** Accuracy of Hamming kNN imputation method at various fractions of random dropout sampled at ∼20% (black) and ∼80% (blue) clonal cell fractions (CCFs). Stem cell population dynamics and long-read barcodes were simulated as in Supplementary Note S7 and sampled at ∼20% and ∼70% CCF, resulting in the ground truth trees show (**b,c**, see methods, Clustering of barcode matrices and calculation of cluster read fractions). **c.** Trees constructed using 100 barcodes with 100 CpGs each with various fractions of CpG dropout (5%, 10%, 20%, and 40%) followed by Hamming kNN imputation of methylation states.

### Supplementary Note 9: Long-read barcode matrices in the PCDH gene cluster

**Table S1.** Long-read matrices coordinates and performance in PD6646 used in Fig. 4 and Fig. S6 (v: variably expressed exon, c: constantly expressed exon)

| Matrix Window | chr:start-end (hg38) | size (kb) | CpGs covered in window | hapA matrix depth | hapA CpGs in matrix | hapB matrix depth | hapB CpGs in matrix |
|---|---|---|---|---|---|---|---|
| pcdha_v1-3 | 5:140784921-140805251 | 20.3 | 706 | 59 | 156 | 67 | 189 |
| pcdha_v2-4 | 5:140793095-140812532 | 19.4 | 683 | 68 | 190 | 68 | 220 |
| pcdha_v3-5 | 5:140805251-140819149 | 13.9 | 252 | 91 | 62 | 80 | 57 |
| pcdha_v4-6 | 5:140812532-140827539 | 15.0 | 242 | 88 | 84 | 74 | 63 |
| pcdha_v5-7 | 5:140819149-140840232 | 21.1 | 714 | 73 | 280 | 61 | 199 |
| pcdha_v6-8.5 | 5:140827539-140842496 | 15.0 | 572 | 80 | 222 | 60 | 180 |
| pcdha_v8.5-10_deletion | 5:140842557-140859330 | 16.8 | 667 | 81 | 234 | (deletion) | (deletion) |
| pcdha_v11-13_1 | 5:140859538-140904808 | 45.3 | 1037 | 81 | 161 | 78 | 218 |
| pcdha_v13+ | 5:140879134-140922061 | 42.9 | 465 | 41 | 122 | 38 | 132 |
| pcdha_c2 | 5:140984494-141018385 | 33.9 | 311 | 64 | 29 | 59 | 34 |
| pcdhb_v1 | 5:141019420-141064247 | 44.8 | 378 | 52 | 58 | 44 | 63 |
| pcdhb_v2-3 | 5:141065070-141115341 | 50.3 | 791 | 56 | 211 | 38 | 149 |
| pcdhb_v4-6 | 5:141112586-141161338 | 48.8 | 842 | 52 | 84 | 60 | 126 |
| pcdhb_v6.5-11 | 5:141152244-141206884 | 54.6 | 1238 | 60 | 205 | 71 | 210 |
| pcdhb_v12-15 | 5:141207549-141252946 | 45.4 | 997 | 70 | 123 | 56 | 161 |
| pcdhb_v15+ | 5:141233413-141258864 | 25.5 | 507 | 78 | 130 | 80 | 111 |
| pcdhga_v1-3 | 5:141327784-141348867 | 21.1 | 505 | 99 | 117 | 72 | 85 |
| pcdhga_v2-4_b1 | 5:141336933-141359156 | 22.2 | 623 | 82 | 207 | 61 | 148 |
| pcdhgb_v2-6 | 5:141358925-141381148 | 22.2 | 686 | 54 | 266 | 53 | 246 |
| pcdhga_v7-b5 | 5:141381097-141403320 | 22.2 | 693 | 63 | 224 | 46 | 223 |
| pcdhga_v9-11_b6-7 | 5:141406930-141424689 | 17.8 | 673 | 78 | 187 | 74 | 113 |

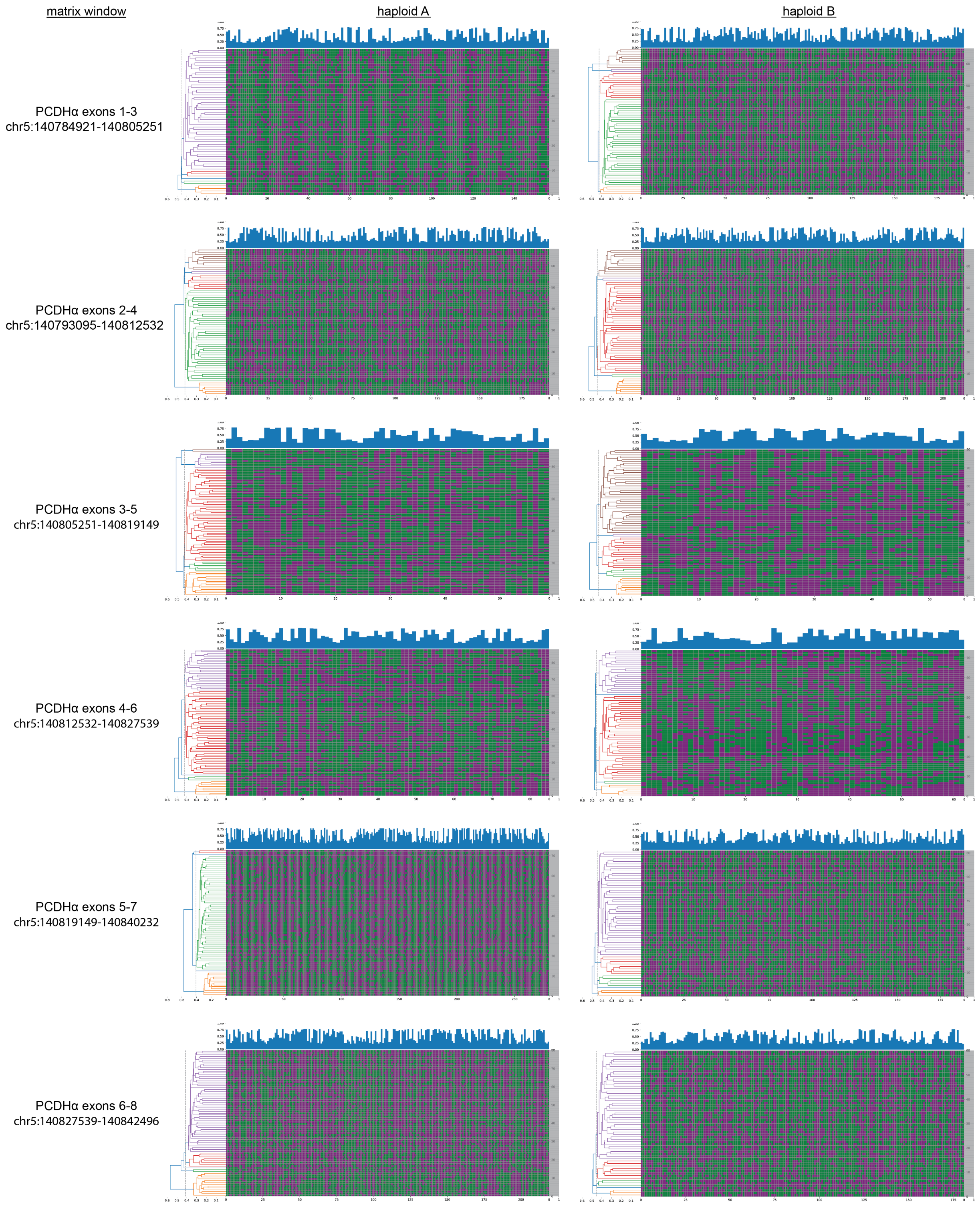


**Fig. S9. Examples of hierarchical clustering of long-read epiallele barcode matrices in the PCDH gene cluster region for patient PD6646.**

### Supplementary Note 10: Additional kmeans clustering of long-read barcodes to quantify cancer cell fractions

We sought to quantify the cancer cell fraction of PD6646 by additional clustering of long-read barcode matrices in the PCDH region, particularly to test approaches that allowed for CpG dropout without imputation. We applied kmeans clustering (k=2 clusters maximum) to each pre-imputation haploid matrix based on the assumption that a cell population with significant clonal diversity would evenly cluster into groups with ~50% with-in group similarity (~50% of CpGs share the same state), or near-random clustering. Whereas, a population with a clonal expansion of <100% clonal cell fraction on top of a wild-type background would result in one cluster with »50% with-in group similarity and one with ~50%, proportional in size to the clonal cell fraction and the remaining wild-type fraction, respectively. We found that kmeans clustering was somewhat better at estimating the DNMT3A clonal cell fraction than hierarchical clustering, it resulted in more orphan clusters that generated bimodal distributions for haploid "B". It was also better at partitioning the more random distributions found in the healthy control.

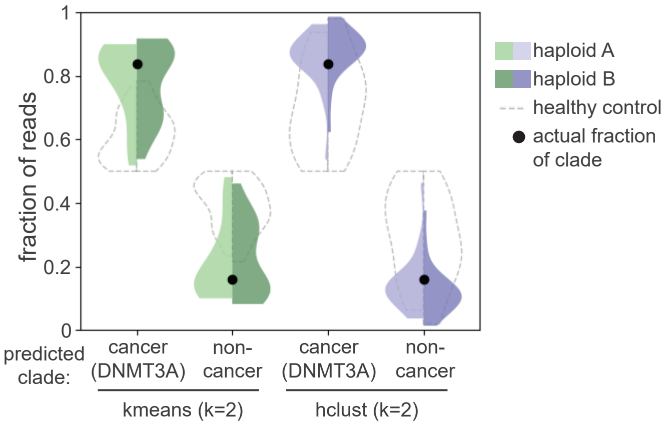


**Fig. S10. Comparison of kmeans and hierarchical clustering methods to infer clade fractions in PD6646 long-read barcode matrices.**

### Supplementary Note 11: Barcode fraction correlation against largest somatic variant size without time-series germline filtering

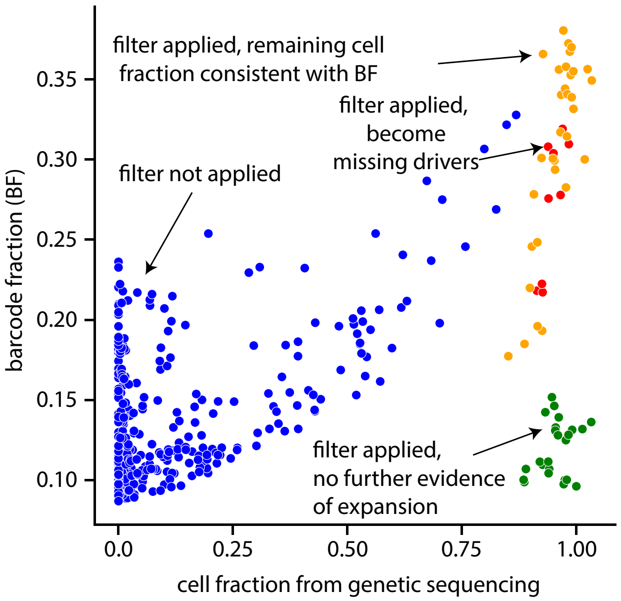


**Fig. S11.** Barcode fraction against variant cell fraction before time-series germline filtering.

### Supplementary Note 12: Evidence of correlations between CpGs

We found that in polyclonal tissue barcodes in which adjacent sites had the same methylation state had higher average read fractions than barcodes in which adjacent sites switch between methylated and unmethylated (supplementary Fig. 12) - there are correlations between nearby sites. This means that in non-clonal samples, certain barcodes are typically larger than would naively be predicted, adding a constant background to statistics that aim to measure clonal size. This lack of independence of epimutations has important implications for using PCDH methylation patterns to build phylogenetic trees, the process of which depends on assumptions about the diversity generation process. We speculate that building a more thorough quantitative understanding of the epimutation process, including of these correlations, may allow us to better reconstruct the lineage history of individuals with clonal expansions.

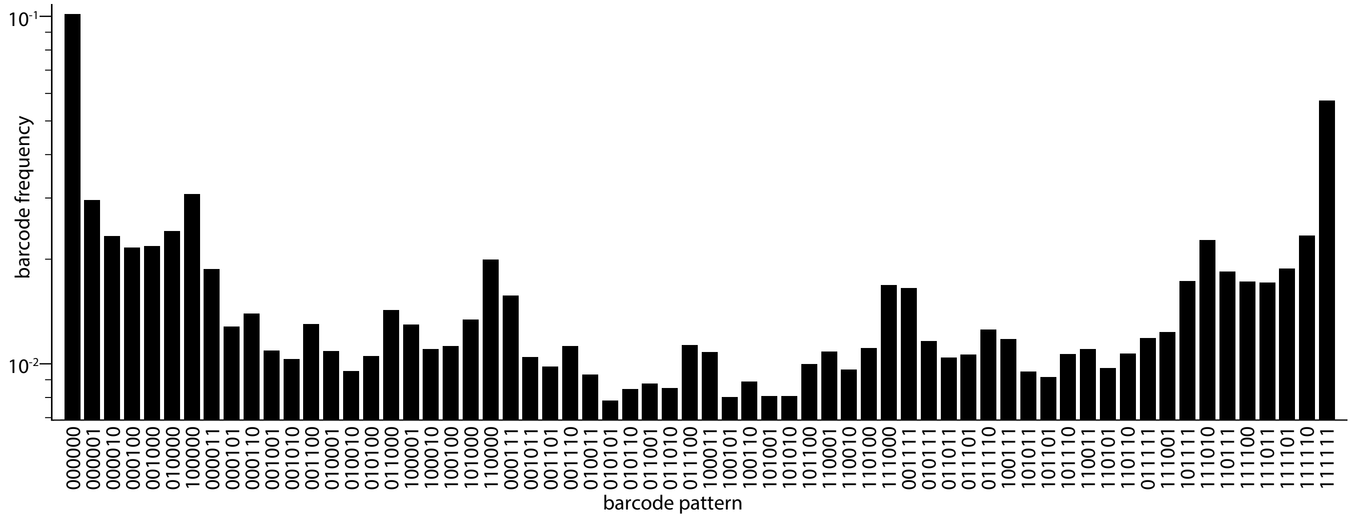


**Fig. S12. Barcode-pattern usage distribution.** The fraction of all 6-mer reads across the protocadherin region which have each of the $2^6 = 64$ possible methylation patterns.

### Supplementary Note 13: Extended data: Longitudinal barcode frequency plots for all samples

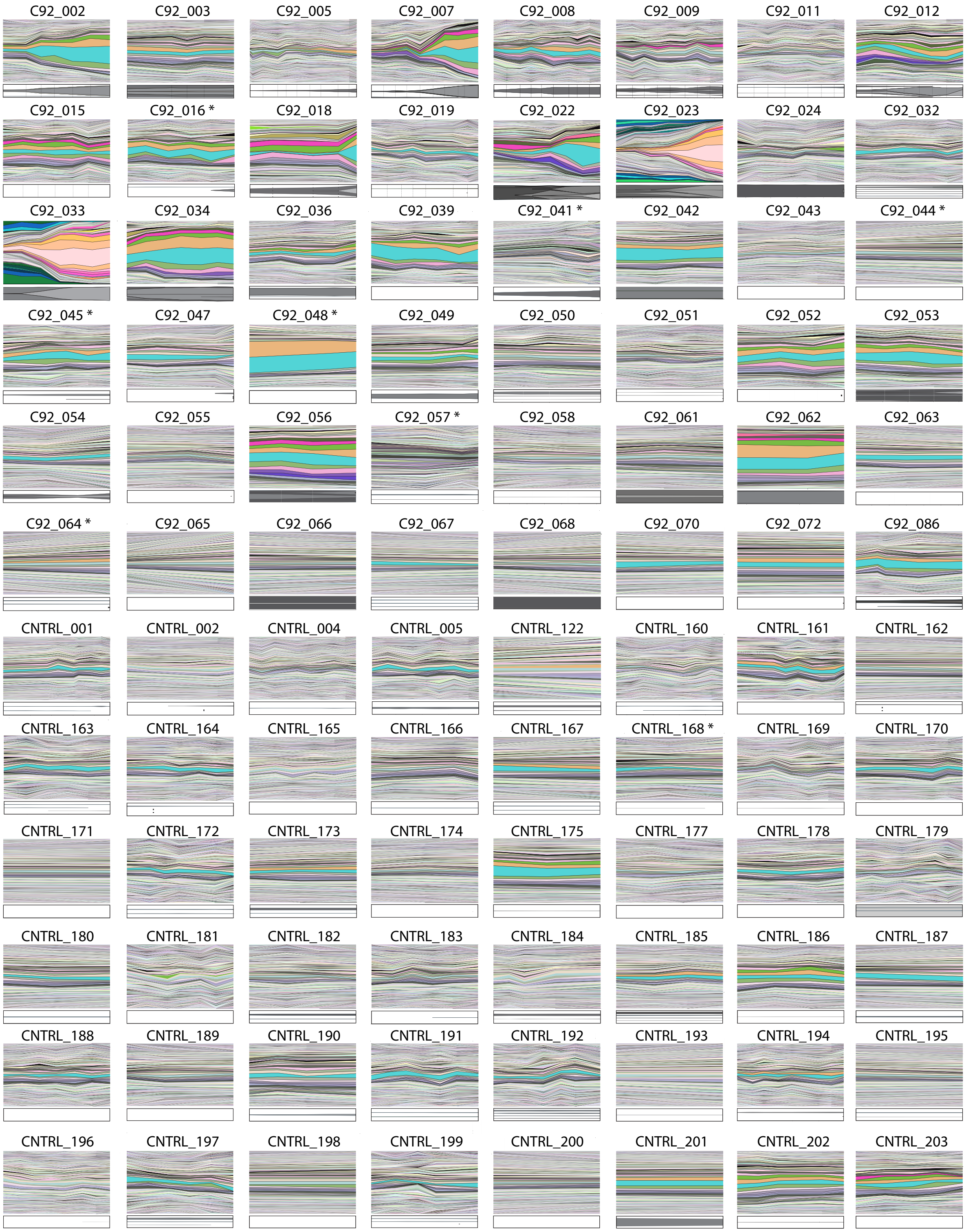


**Fig. S13. Longitudinal barcode frequency plots for all samples.** Longitudinal plots for all samples at the ten-mer starting at hg19 chr5:140256590 (the same as used in Fig. 1). For eight individuals (starred), there was insufficient depth at some timepoint at this locus, and so the first locus which had sufficient samples in all samples was used - if there was no such locus, the first locus which maximised the number of samples with sufficient coverage was used, and the samples without sufficient depth were excluded.

### Supplementary Note 14: Barcodes that expand at a given site are chosen randomly

The barcodes which expanded at a given locus during a clonal sweep were different across individuals (supplementary Fig. 14a), consistent with the hypothesis that these patterns act as neutral lineage markers. To quantify this we found the average Hamming distance between the two expanded barcodes across all non-constant PCDH loci. If barcodes were chosen at random, the expected value of this quantity would be $l((m_1 + m_2) - 2m_1m_2)$, where $m_1$ and $m_2$ are the average methylations of two barcodes respectively and $l$ is the barcode length (we use $l = 10$). Both on the two haploids of one individual (supplementary Fig. 14b) and across individuals (supplementary Fig. 14c), the established barcodes are no more closely related than would be expected by chance.

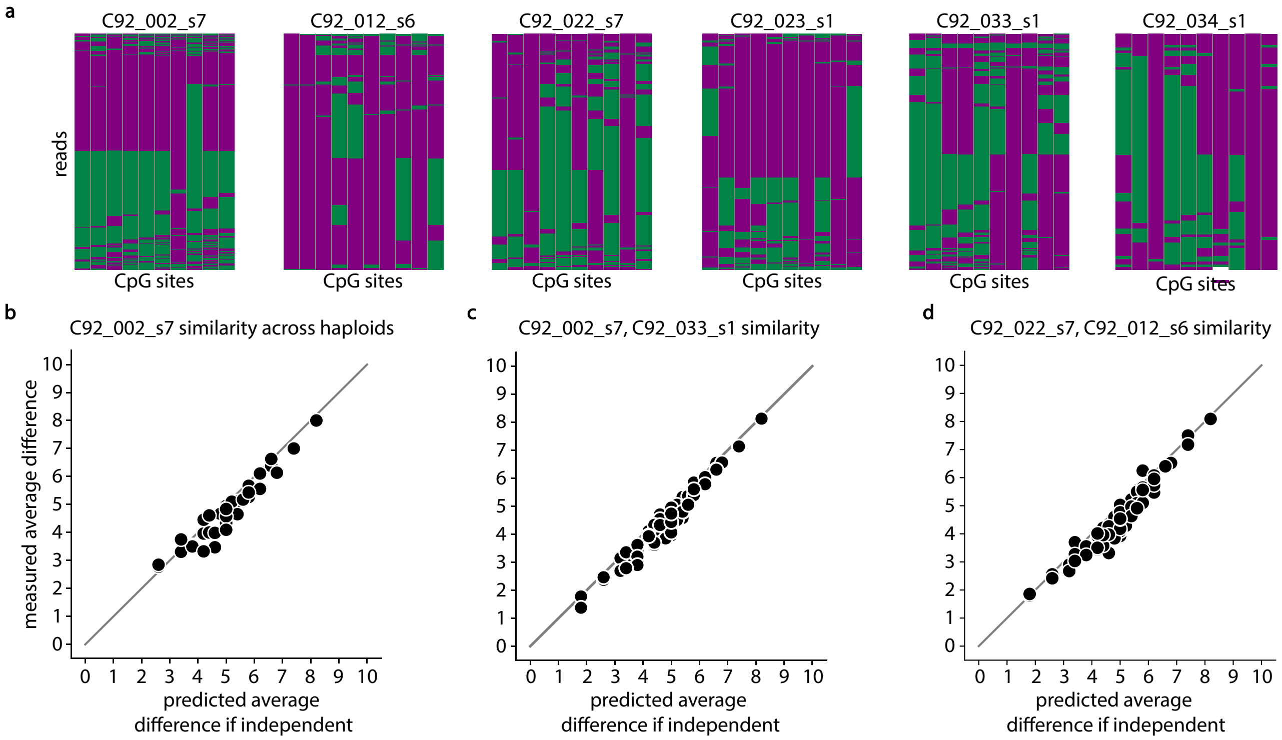


**Fig. S14. Barcodes that expand at a given locus are chosen randomly. a.** Example pileups at a locus in PCDH (hg19 chr5:140167118) **b.** The average Hamming distance between the estimated founding barcodes on each chromosome for C92_002_s7, compared to the difference expected if barcode patterns across haploids were completely unrelated. **c.** The average Hamming distance between founding barcodes at the same loci in C92_002_s7 and C92_033_s1, compared to the theoretical expectation if barcodes were independent. **d.** The same for C92_022_s7 and C92_012_s6.

### Supplementary Note 15: Barcode statistics give lower bound on solid tissue sample purity

We reasoned that the BF in solid tissue samples should give a lower bound to tumour purity, which is analogous to the cell fraction in figure 2. Comparing this statistic with Battenburg sample-purity estimates produces the expected relationship in both prostate tumour samples (supplementary Fig. S15 a) and clear cell renal carcinoma samples (supplementary Fig. S15 b).

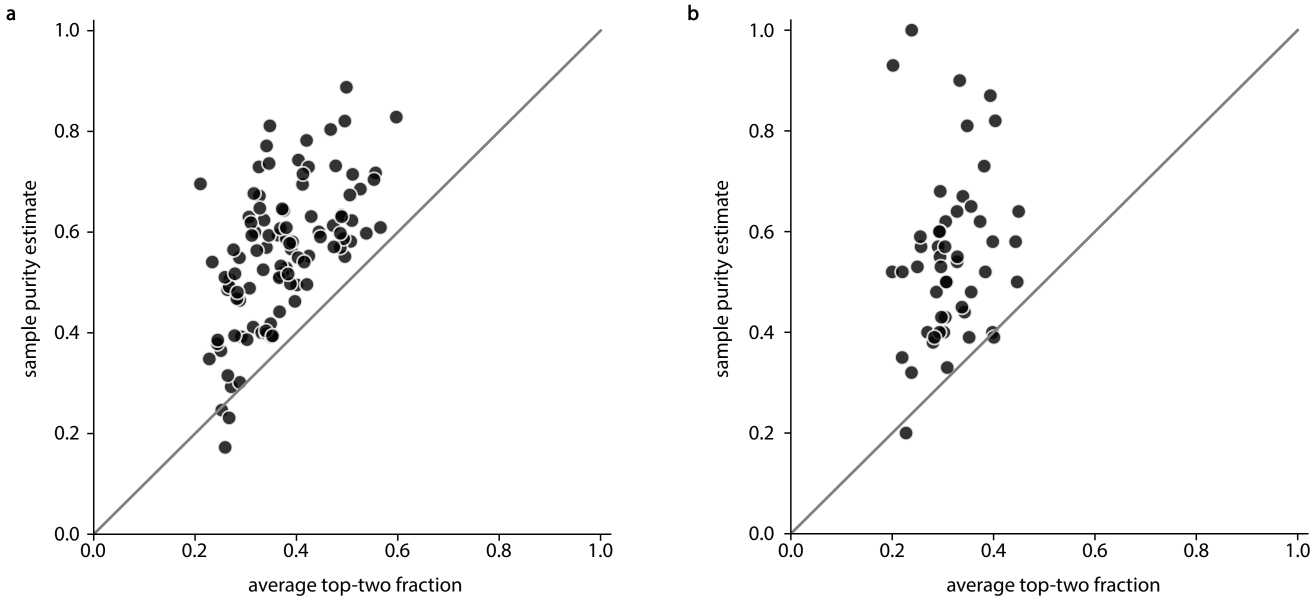


**Fig. S15. Methylation barcode statistics give lower bounds for solid-tissue sample purity. a.** Scatter plot showing the relationship between the 'top-two' barcode statistic (x-axis) and sample purity estimate (y-axis) for prostate tumour samples. **b.** The same for renal cancer samples. Grey lines indicate $y = x$.

### Supplementary Note 16: Extended data: Longitudinal barcode frequency plots for all non-overlapping ‘good-eloci’ for C92_002

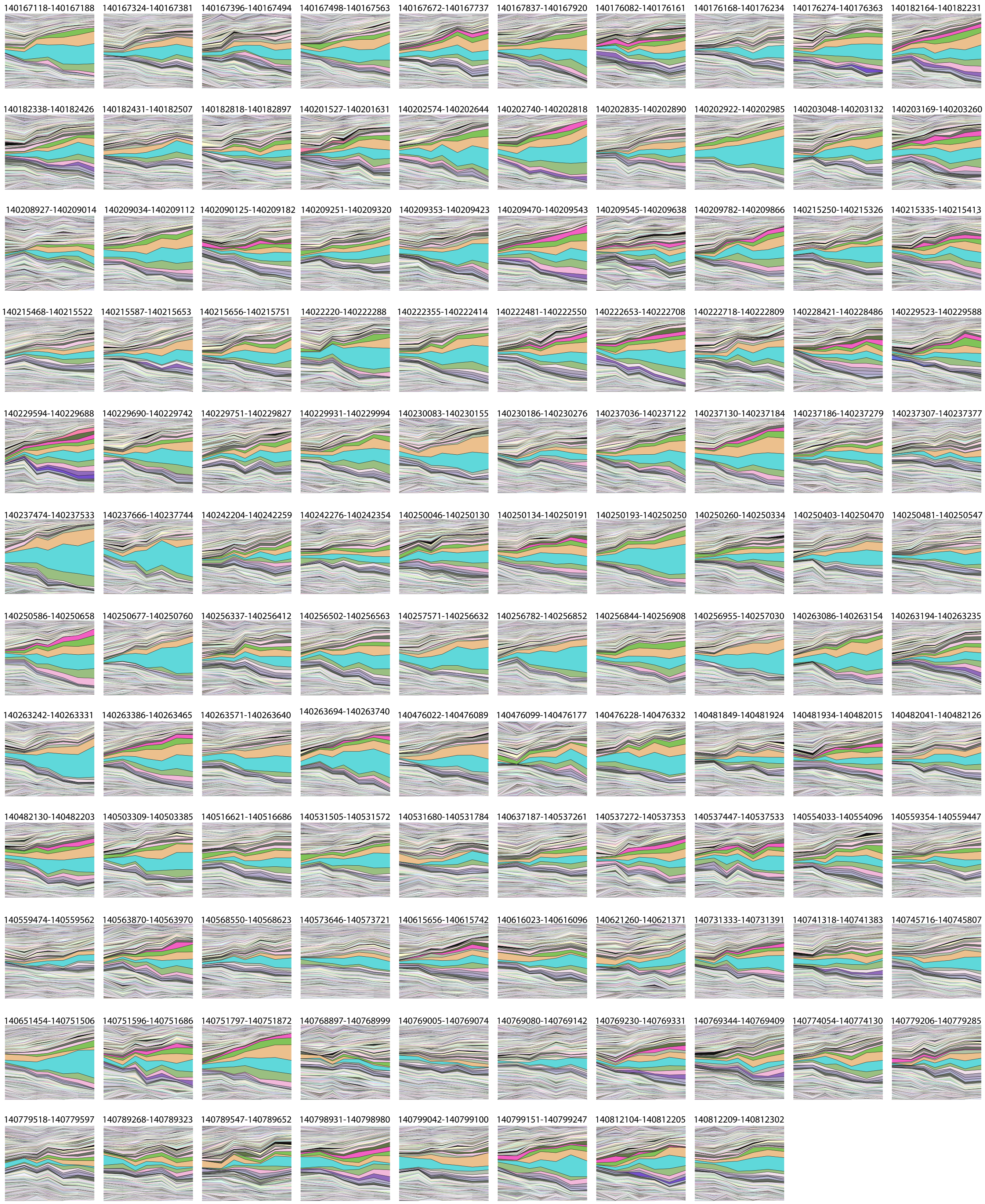


**Fig. S16.** Barcode-frequency plots for all non-overlapping ‘good’ eloci for C92_002.

### Supplementary Note 17: Extended data: Longitudinal barcode frequency plots for all non-overlapping ‘good-eloci’ for C92_033

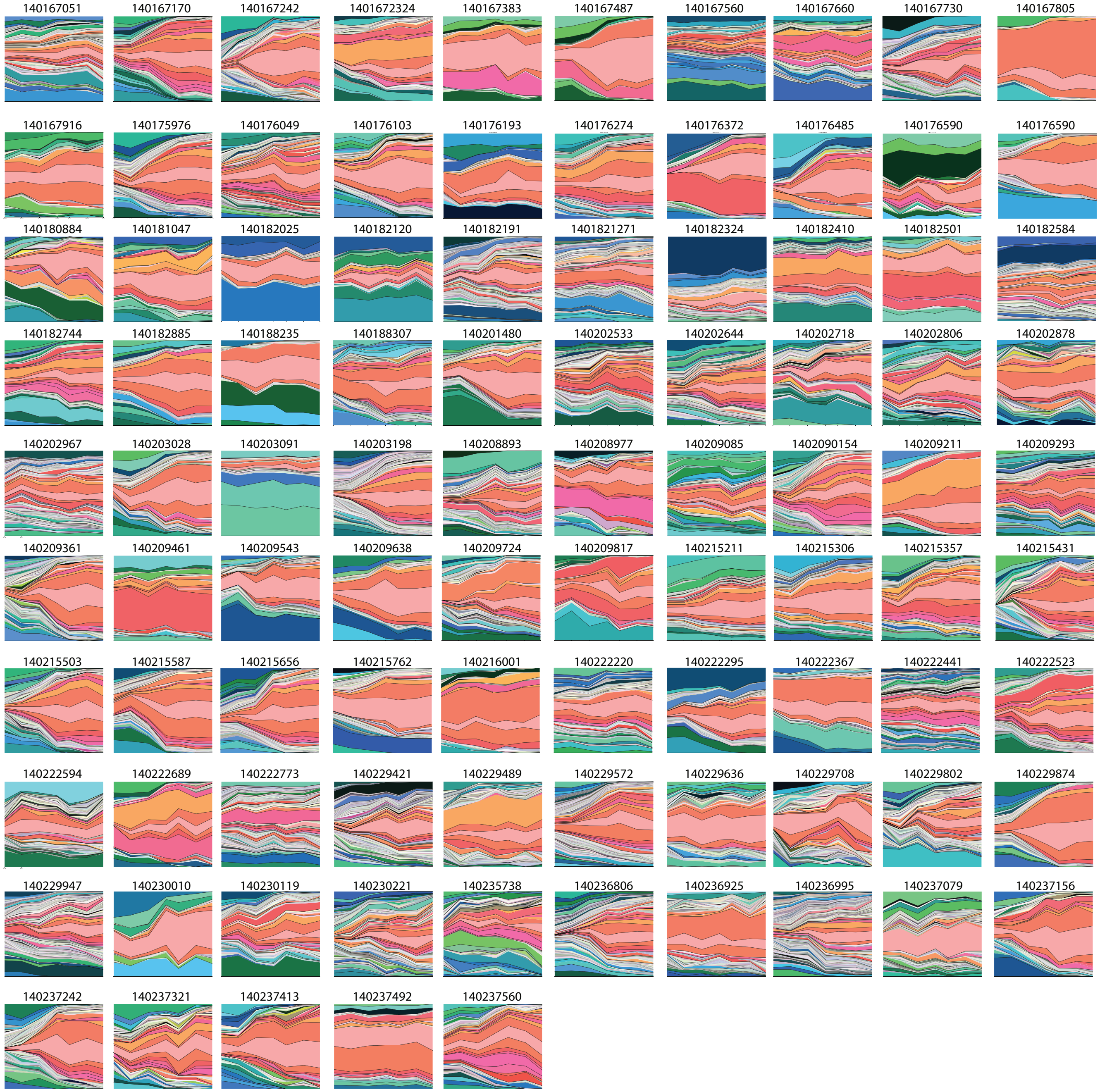


**Fig. S17.** Barcode-frequency plots for all non-overlapping ‘good’ eloci for C92_033.

### Supplementary Note 18: Extended data: Entropy in individuals with and without large clonal sweeps across panel

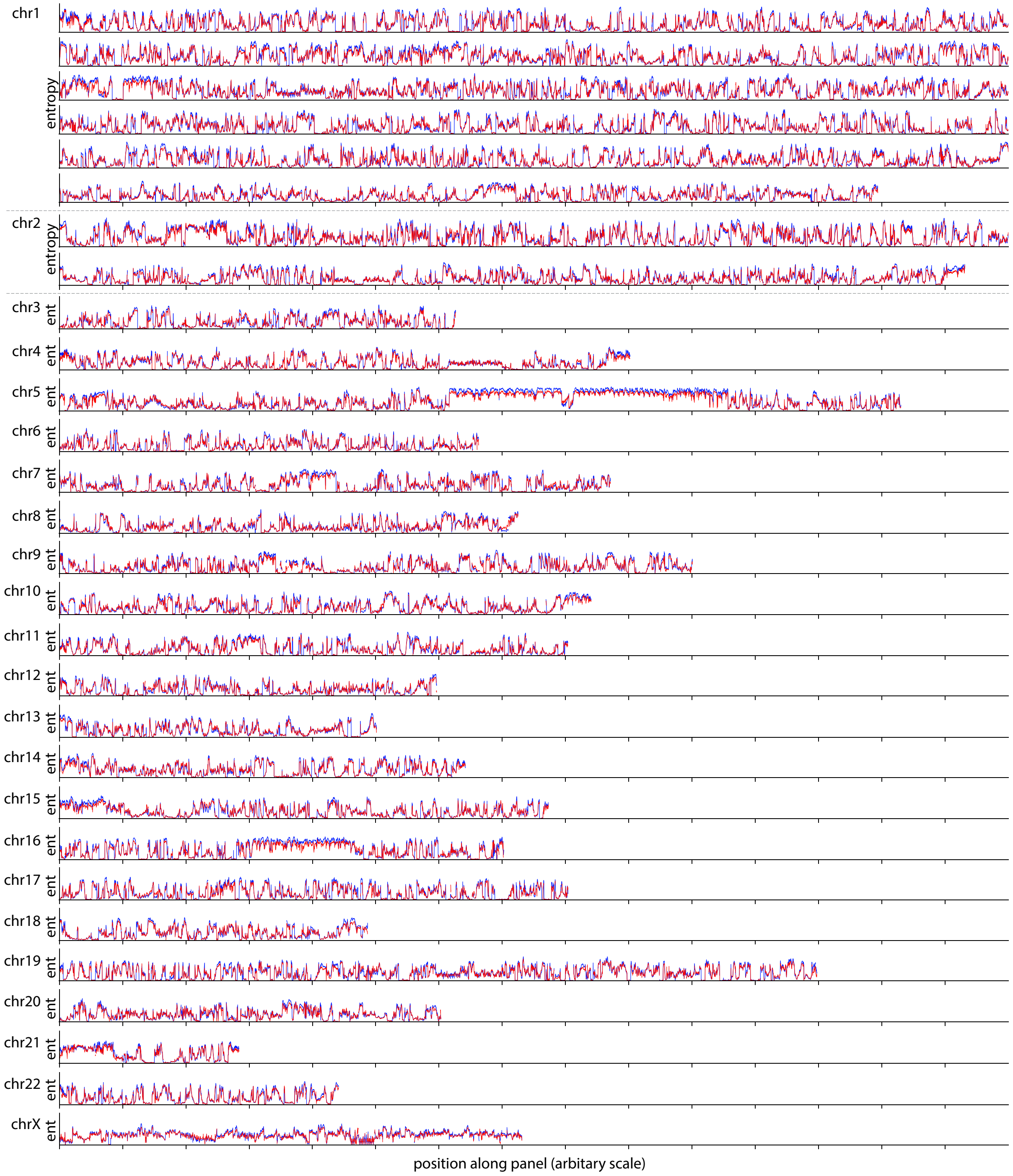


**Fig. S18.** 10-mer methylation entropy averaged across individuals with large clonal sweeps ($> 80\%$ cell fraction, red) and without detected large clonal sweeps ($< 20\%$ cell fraction, blue). Ticks on x axis indicate 1000 (possibly overlapping) ten-mers.

### Supplementary Note 19: Entropy across PCDH in lung cancer samples

We considered the entropy across the protocadherin region in a set of lung cancer samples (small cell lung cancers - SCLC, $n = 18$ and lung adenocarcinomas - LUAD, $n = 7$)[59]. The protocadherin region had a remarkably similar entropy profile in these samples as was observed in blood, kidney, prostate, and bladder samples. The lack of sequencing data from healthy lung precluded further analysis.

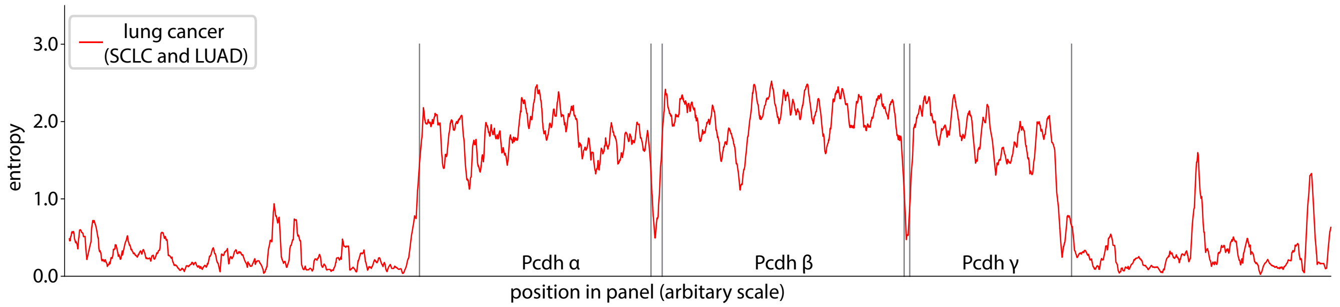


**Fig. S19.** Average entropy profile across the protocadherin region in lung cancer samples.
